# NOTCH1 S2513 is critical for the regulation of NICD levels impacting the segmentation clock in hiPSC-derived PSM cells and somitoids

**DOI:** 10.1101/2024.12.10.627712

**Authors:** Hedda A. Meijer, Adam Hetherington, Sara Johnson, Rosie L. Gallagher, Izzah N. Hussein, Jess M. Rae, Tomas E.J.C. Noordzij, Margarita Kalamara, Thomas J. Macartney, Lindsay Davidson, David M.A. Martin, Paul Davies, Katharina F. Sonnen, Philip Murray, J. Kim Dale

**Affiliations:** Division of Molecular Cell and Developmental Biology, School of Life Sciences, University of Dundee, Dow Street, Dundee DD1 5EH, United Kingdom; Mathematics, School of Science and Engineering, University of Dundee, Dow Street, Dundee DD1 4HN, United Kingdom; Hubrecht Institute, KNAW (Royal Netherlands Academy of Arts and Sciences), University Medical Center Utrecht, 3584 CT Utrecht, the Netherlands; Human Pluripotent Stem Cell Facility, School of Life Sciences, University of Dundee, Dow Street, Dundee DD1 5EH, United Kingdom; MRC-PPU, School of Life Sciences, University of Dundee, Dow Street, Dundee DD1 5EH, United Kingdom; D’Arcy Thompson Unit, School of Life Sciences, University of Dundee, Dow Street, Dundee DD1 5EH, United Kingdom

**Keywords:** NOTCH1, segmentation clock, somitogenesis, hiPSC, PSM, somitoid

## Abstract

The segmentation clock is a molecular oscillator that regulates the timing of somite formation in the developing vertebrate embryo. NOTCH signalling is one of the key pathways required for proper functioning of the segmentation clock. Aberrant NOTCH signalling results in developmental abnormalities such as congenital scoliosis as well as diseases such as T-cell acute lymphoblastic lymphoma (T-ALL). In this study we analyse the effects of a mutation detected in T-ALL patients on somitogenesis using human iPS derived PSM cells and somitoids. Mutation of NOTCH1 Serine 2513 into Alanine compromises the interaction of Notch intracellular domain (NICD) with the F-box protein FBXW7 and consequently increases NICD stability and NICD levels in PSM cells. Moreover, the mutation impairs several aspects of clock gene oscillations and restricts the ability of somitoids to polarise, elongate and form paired somites. The data suggest a mechanism by which post-translational modification of a key segmentation clock component plays a crucial role in vertebrate axis segmentation.

## Introduction

Somitogenesis is a process that occurs early in the development of the vertebrate embryo. It eventually gives rise to the bones and muscles of the vertebrate skeleton and some of the dermis. Malfunctioning somitogenesis can result in musculoskeletal deformities, leading to conditions such as Spondylocostal Dysostosis (Nóbrega et al, 2021). During somitogenesis blocks of cells, known as somites, segment off the anterior end of the presomitic mesoderm (PSM) at periodic time intervals. Concurrently, the PSM is replenished with cells from the primitive streak and later the tailbud (Hubaud and Pourquié, 2014). The timing of somite formation is regulated by the periodic expression of segmentation clock genes, many of which exhibit posterior to anterior expression waves. As cells become anteriorly placed in the PSM, they undergo a mesenchymal to epithelial transition (MET) that ultimately leads to the formation of discrete somite borders (Cooke and Zeeman, 1976; reviewed by McDaniel et al, 2024). The periodicity of both the segmentation clock and somite formation are tightly correlated and highly species specific, varying from 30 min in zebrafish to 5 hours in human embryos (reviewed by Carraco et al., 2022). This timing is determined by factors such as processing delays and the half-lives of unstable clock gene mRNAs and proteins (Lazaro et al., 2023).

Segmentation clock gene expression is driven by three different pathways (NOTCH, WNT and FGF) that are interlinked (Dequéant et al, 2006) and eventually result in pulses of ppERK which in turn enables segment formation (Simsek et al., 2023). NOTCH is the key signalling pathway for the coordination of clock gene expression (Jiang et al., 2000). In canonical *in trans* NOTCH signalling, the NOTCH receptor, a transmembrane protein in the signal-receiving cell, interacts with transmembrane proteins DELTA or JAGGED in signal-sending cells. This results in several cleavage events, mediated by the ADAM17 and gamma secretase proteases, and the eventual release of NICD. NICD then translocates to the nucleus where it activates NOTCH target genes by complexing with the RBPJ/CSL and MAML transcription factors (reviewed by Kopan and Ilagan, 2009; Ramesh and Chu, 2024). The NOTCH signalling pathway is one of the key pathways critical for accurate clock gene expression profiles in PSM cells (Jiang et al, 2000; Ferjentsik et al, 2009; Loureiro et al., 2024). Moreover, linkage analysis has detected mutations in several NOTCH target genes in patients with Spondylocostal Dysostosis (Nóbrega et al., 2021).

Efficient NICD degradation is necessary for Delta-Notch signalling to exert a short-lived effect on target gene expression. NICD is highly labile, and its degradation is mediated via phosphorylation (reviewed by Borggrefe et al., 2016) and the subsequent recruitment of E3 ubiquitin ligases (reviewed by Dutta et al., 2022). SCF E3 ligase, with substrate recognition component FBXW7, is the predominant E3 ligase involved in NICD ubiquitination and subsequent proteasomal degradation. The NICD PEST domain contains an FBXW7 degron that includes Serine 2513 (Thompson et al., 2007). This residue has been shown to be required for NICD-FBXW7 interaction in HEK293 cells (Carrieri et al., 2019) and has also been identified as one of the mutated residues in T-ALL patients (Bonn et al., 2013). T-ALL is a cancer characterised by faulty NICD degradation and more than 60% of patients carry *NOTCH1* (Weng et al., 2004) or *FBXW7* mutations (Yeh et al., 2016). T-ALL patients carrying *NOTCH1* mutations show increased NICD levels in primary leukaemia cells (Zuurbier et al., 2010) as well as increased mRNA expression of NOTCH1 targets such as *HES1* (Zuurbier et al., 2010; Fogelstrand et al., 2014). Although there is some understanding of the processes that regulate NICD stability in contexts such as T-ALL, very little is known in the context of embryogenesis. It has been shown that *NOTCH1* and *DELTAlike1* mRNA, NICD protein, as well as a variety of NOTCH1 target genes are dynamically expressed in the PSM of chick and mouse embryos (reviewed by Ramesh and Chu, 2024). The periodicity of this dynamic expression is regulated by the positive and negative feedback loops of unstable regulators (Kageyama et al., 2018). NICD itself is one of those key regulators.

Recent protocols for the differentiation of human induced pluripotent stem cells (hiPSC) into PSM cells (Diaz-Cuadros et al., 2020) enable the production of large numbers of PSM cells *in vitro*. Using a modified YFP reporter ACHILLES under control of the promoter of the clock gene *HES7 (HES7-ACHILLES*), it has been shown that the segmentation clock oscillates with a period of approximately 5 hours (Diaz-Cuadros et al., 2020), requiring timely production and degradation of the oscillating mRNAs and proteins encoded by the clock genes. Protocols for the generation of hiPSC-derived organoids that form somite-like segments, termed somitoids, have also been recently developed (Sanaki-Matsumiya et al., 2022; Yamanaka et al., 2023; Miao et al 2023). Both the PSM differentiation and somitoid protocols now allow for the analysis of mutations found in patients to be investigated using *in vitro* generated human model systems. Using gene editing of hiPSCs and the generation of human PSM cells and somitoids, here we provide data to show that, in PSM cells, NICD stability is regulated via FBXW7 interaction with NICD S2513 in the PEST domain. We also demonstrate that mutation of this Serine residue into an Alanine disrupts FBXW7 mediated regulation of NICD levels, resulting in aberrant NICD and clock gene expression as well as defects in somitoid development.

## Results

### FBXW7 is required for the regulation of NICD levels in iPSC-derived PSM cells

NICD is a critical activator of clock gene expression in the PSM. The tight regulation of the dynamic aspects of NICD activation and turnover are therefore essential for the timing of the segmentation clock. To establish the half-life of NICD in PSM cells, hiPSCs were differentiated into PSM cells and treated with the gamma secretase inhibitor LY411575 to block the release of new NICD from the plasma membrane. A subsequent time course enabled inference of the NICD half-life by western blotting. Hence, we determined that the NICD half-life in Wibj2 (https://www.hipsci.org) hiPSC-derived PSM cells is approximately two hours (Fig 1A). Notably, this value is less than half of the duration of the segmentation clock cycle in human cells (Diaz-Cuadros et al, 2020; Carraco et al, 2022).

**Figure 1.**
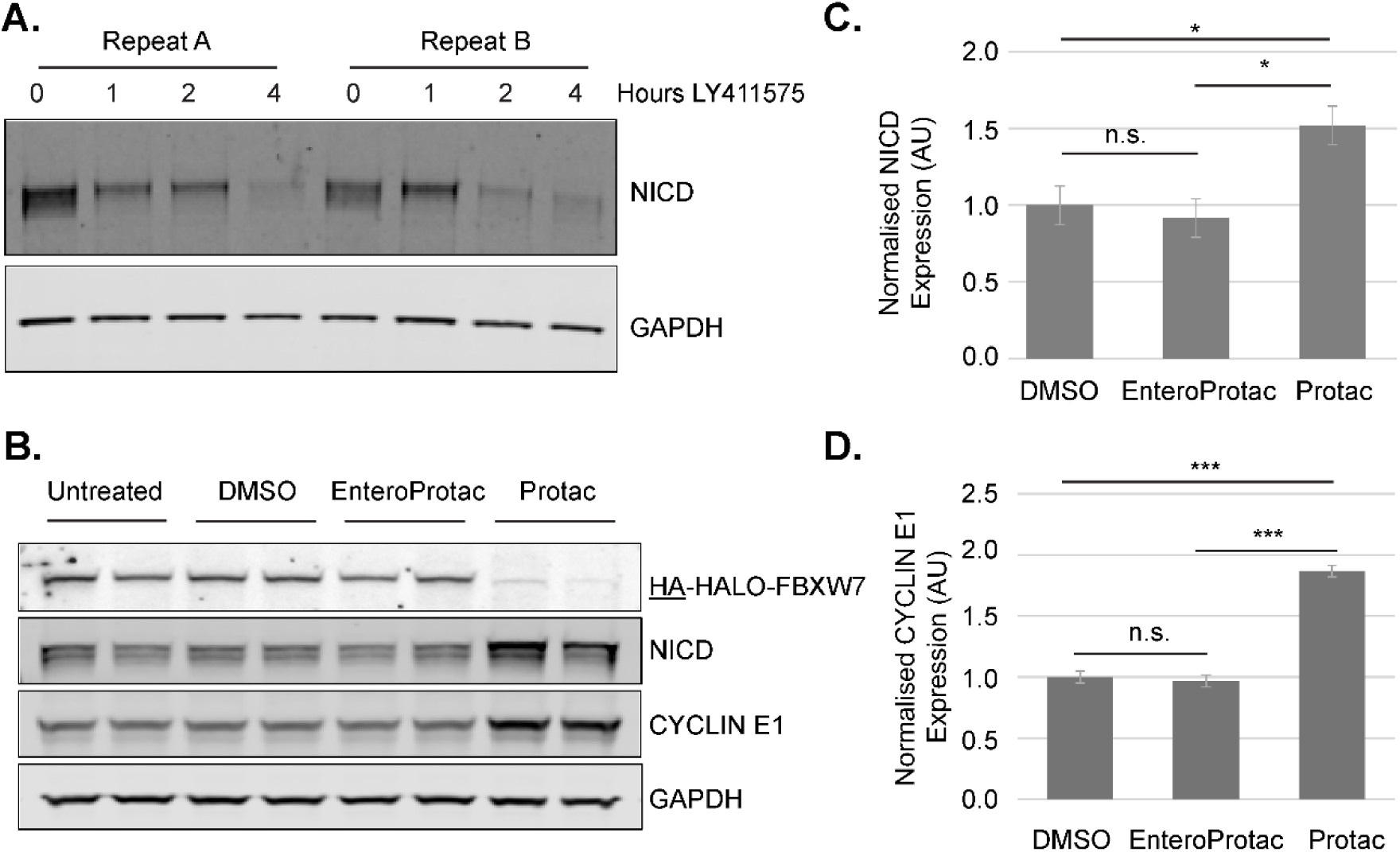
FBXW7 is required for the regulation of NICD levels in PSM. A) Wibj2 PSM cells were treated with LY411575 and harvested several hours after LY addition as indicated and analysed by WB. Representative experiment is shown. NICD levels from were quantified for three biological repeats (two technical repeats each), normalised to GAPDH and NICD half-life was calculated (1.83 hours). B) *HA-HALO-FBXW7* PSM cells were treated with PROTAC / enteroPROTAC / DMSO for 12 hours. Analysis of expression levels of FBXW7 targets by WB. Representative experiment is shown. C/D) Quantification of four biological repeats (two technical repeats each). The intensity of NICD and CYCLIN E1 bands were normalised to ACTB and values are displayed relative to vehicle-only DMSO treated values (mean +/- s.e.m.). The ablation of HA-HALO-FBXW7 with PROTAC causes a significant increase in expression of NICD, compared to DMSO (t=2.935, df=6, p=0.0261) and enteroPROTAC controls (t=3.394, df=6, p=0.0146) and also of CYCLIN E1 vs DMSO (t=12.635, df=6, p<0.0001) and vs enteroPROTAC (t=13.097, df=6, p<0.0001). No significant differences were observed when comparing DMSO and enteroProtac samples.

To establish whether FBXW7, one of the subunits of the SCF (SKP1-CULLIN-FBOX) E3 ubiquitin ligase complex, is involved in the regulation of NICD degradation in iPS-derived PSM cells, as reported in other cellular contexts (Dutta et al., 2022), we assayed NICD levels upon FBXW7 perturbation. Using CRISPR-CAS9 gene editing, hemagglutinin (HA) and HALO tags were added to the endogenous *FBXW7* locus (*HA-HALO-FBXW7*, Fig S1) of *HES7*-ACHILLES (*HES7A*) hiPSCs (Diaz-Cuadros et al., 2020). The HALO tag enables the targeting of FBXW7 for fast degradation using PROTACs, protein degraders that activate the E3 ubiquitin ligase machinery (Webb et al., 2022). Sequencing of both strands of the DNA, including and surrounding the knock in site, confirmed the homozygous addition of the tags (data not shown). Verification of this modified hiPSC cell line confirmed the expression of pluripotency markers (Fig S2A-D, Table S1) and the ability to differentiate into all three germ layers (Table S2). Moreover, the introduction of the HA and HALO tags did not affect the ability of these cells to differentiate into PSM cells (Fig S3A) or the expression of the FBXW7 targets NICD and CYCLIN E1 (Fig S4A-B). Furthermore, when the HALO tag was targeted by PROTAC following exposure for 6h FBXW7 was, as expected, efficiently degraded in hiPSCs (Fig S4C). The treatment of iPS-derived PSM cells with PROTAC also resulted in the degradation of FBXW7 and subsequent increased expression of the FBXW7 targets NICD and CYCLIN E1 (Fig 1B-D). No effect was observed with the enteroPROTAC which is an inactive form of PROTAC. This demonstrates for the first time that FBXW7 regulates NICD levels in PSM cells.

### Serine 2513 (S2513) in the NICD PEST domain is critical for the regulation of NICD half-life in PSM cells

The NICD PEST domain has been reported to regulate NICD stability (Thompson et al., 2007). To determine if the FBXW7-NICD interaction is mediated via one of these residues in the NICD PEST domain (S2513), the *HA-HALO-FBXW7* hiPSCs were further modified using CRISPR-CAS9 to mutate NOTCH1 Serine 2513 into an Alanine, thus preventing the phosphorylation of S2513. Moreover, an mCHERRY tag was added to both the wild type (WT) endogenous *NOTCH1* locus and the S2513A mutant *NOTCH1* locus to enable the FACS sorting of modified clones (WT *NOTCH1* and S2513A *NOTCH1*, Fig S1). Structural analysis of the fusion proteins (using AlphaFold, https://alphafold.com) suggests that the modifications are unlikely to affect the structure of NICD (data not shown). Sequencing of both strands of the DNA, including and surrounding the knock in site, confirmed the homozygous addition of the tags and the mutation (data not shown). Verification of these two new cell lines confirmed pluripotency marker expression (Fig S2E-H, Table S1) as well as the ability to differentiate into the three different germ layers (Fig S3B, Table S2). Notably, although differentiation into PSM was not affected by the S2513A mutation, it did strongly perturb differentiation into neuroectoderm, as evidenced by a clear reduction in PAX6 expression compared to wild type cells exposed to the neuroectoderm differentiation assay (Fig S3C). These data suggest that increasing NICD levels in a bipotential progenitor cell population biases differentiation to a mesodermal rather than a neuronal fate, as has been recently reported (Cooper et al., 2023).

To determine the effect of the S2513 mutation in PSM cells, NICD levels in WT and mutated cell lines were assayed using western blot analysis. The S2513A mutation resulted in significantly increased NICD expression levels with only minor non-significant effects on cleaved NOTCH1 expression levels (Fig 2A-B). Full length NOTCH1 levels were not determined as neither the NOTCH1 nor the mCHERRY antibody generated a reliable signal for full length NOTCH1 (data not shown). The lower NICD band (*), which is detected by the NICD antibody but not by mCHERRY WB (data not shown), is predicted to be NICD plus linker and a small part of mCHERRY. To investigate whether the increased NICD levels in S2513 cells were a consequence of stabilised NICD, WT and S2513A *NOTCH1* PSM cells were treated with LY411575 (thus blocking release of cleaved NICD) in order to determine the NICD half-life (Fig 2C-D). Whilst NICD half-life in the WT *NOTCH1* PSM cells was similar to that observed in Wibj2 PSM cells (∼ 2 hours; Fig 1A), it was significantly increased in S2513A *NOTCH1* PSM cells (∼5 hours). These data confirm that NOTCH1 S2513 regulates NICD stability and NICD levels in PSM cells.

**Figure 2.**
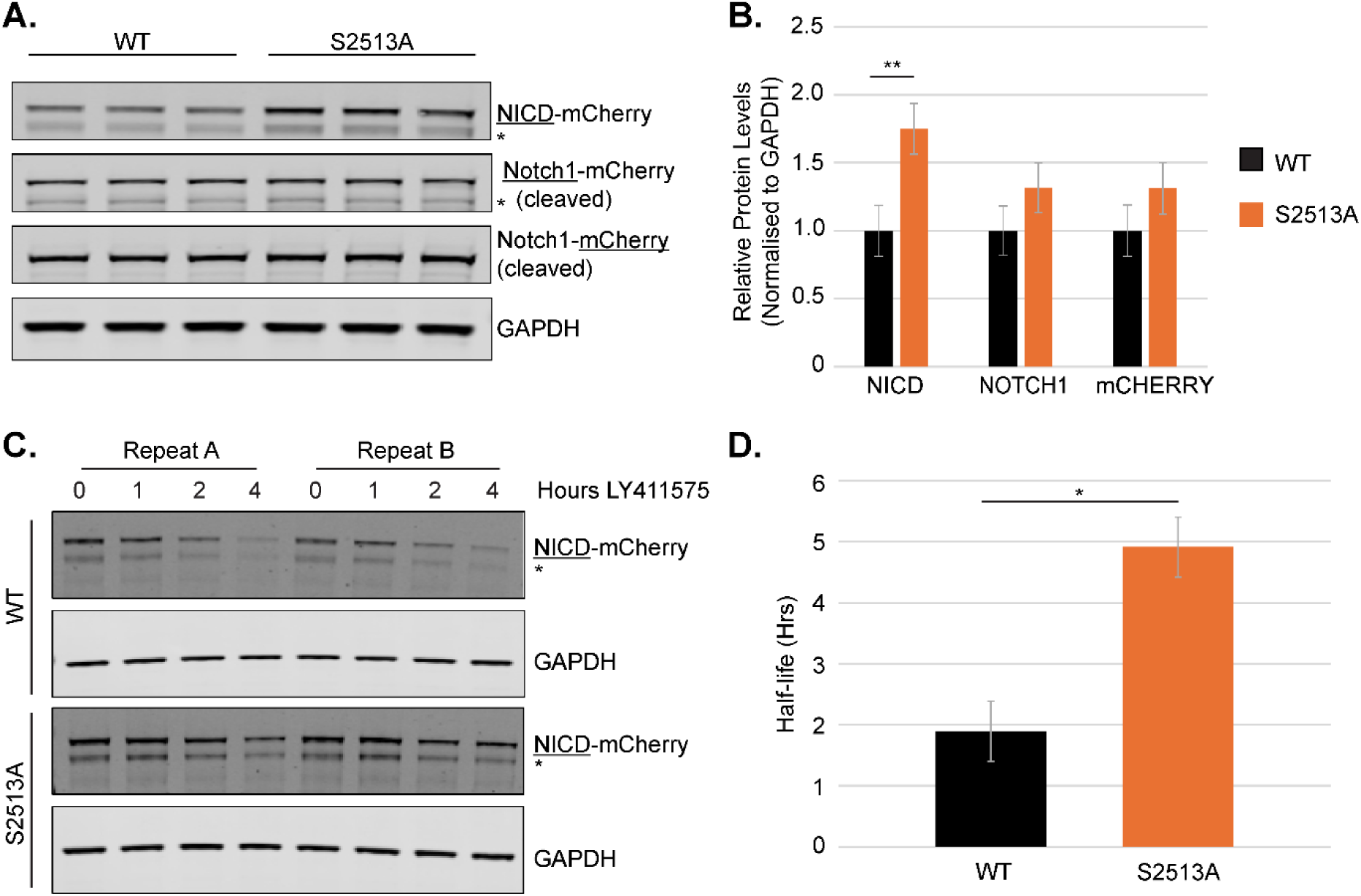
Modification of NOTCH1 S2513 disrupts the regulation of NICD protein levels in the PSM. A) WB analysis of NOTCH1 and NICD expression levels in WT and S2513A *NOTCH1* PSM cells. Representative experiment is shown. B) Quantification of three biological repeats (three technical repeats each). Protein levels were normalised to GAPDH (mean +/- s.e.m.). NICD levels are increased 1.75x in S2513A *NOTCH1* PSM cells (t=3.194, df=13.5 p=0.0067) with no significant effect on cleaved NOTCH1 (t=1.461, df=13.4, p=0.1671) or mCHERRY (t=1.391, df=13.3, p=0.1869) levels. C) WT and S2513A *NOTCH1* PSM cells were treated with LY411575 and harvested several hours after LY411575 addition as indicated and analysed by WB. Representative experiment is shown. NICD levels from C) were quantified for three biological repeats (two technical repeats each), and normalised to GAPDH. At all timepoints the normalised amount of remaining protein is different between the two cell lines (1 hour: t=2.611, df 38.1, p=0.0128; 2 hours: t=2.333, df=38.1, p=0.0250; 4 hours: t=4.123, df=38.1, p=0.0002). D) The half-life of NICD measured at the 4-hour time point in S2513A *NOTCH1* PSM cells is significantly increased by ∼3 hours compared to WT *NOTCH1* PSM cells (1.89 hours for WT and 4.91 hours for S2513A *NOTCH1* PSM cells; t=4.333, df=4, p=0.0123). *indicates NICD lacking most of mCHERRY.

To determine whether the S2513 residue is required for the NICD-FBXW7 interaction, *HA-HALO-FBXW7*, WT and S2513A *NOTCH1* PSM cell lysates were subjected to mCHERRY immunoprecipitation (IP) (Fig 3A). The *HA-HALO-FBXW7* cell line serves as a negative control for the IP as it does not contain an mCHERRY tag. A significantly reduced interaction between NICD and HA-HALO-FBXW7 was observed in the S2513A *NOTCH1* PSM cells. This is despite the increased NICD levels in total cell lysate and subsequently the precipitation of more NICD in the S2513A *NOTCH1* cell line compared to the WT *NOTCH1* cell line (Fig 3A-B). To establish the relevance of the Serine 2513 residue for FBXW7 mediated degradation of NICD, *NOTCH1* WT and S2513A PSM cells were treated with PROTAC. As expected, exposure to PROTAC resulted in significantly increased levels of the FBXW7 target CYCLIN E1 in both WT and S2513A *NOTCH1* PSM cells (Fig 3C-E). In contrast, whereas NICD levels were significantly higher in WT *NOTCH1* PSM cells following exposure to PROTAC (Fig 3C/F), there was no significant difference in NICD levels in S2513A *NOTCH1* PSM cells after PROTAC treatment, showing a lack of response of NICD carrying the S2513 point mutation to the removal of FBXW7 (Fig 3C/G). Together, these data suggest that S2513 in the NICD PEST domain is required for efficient FBXW7-mediated NICD degradation in PSM cells.

**Figure 3.**
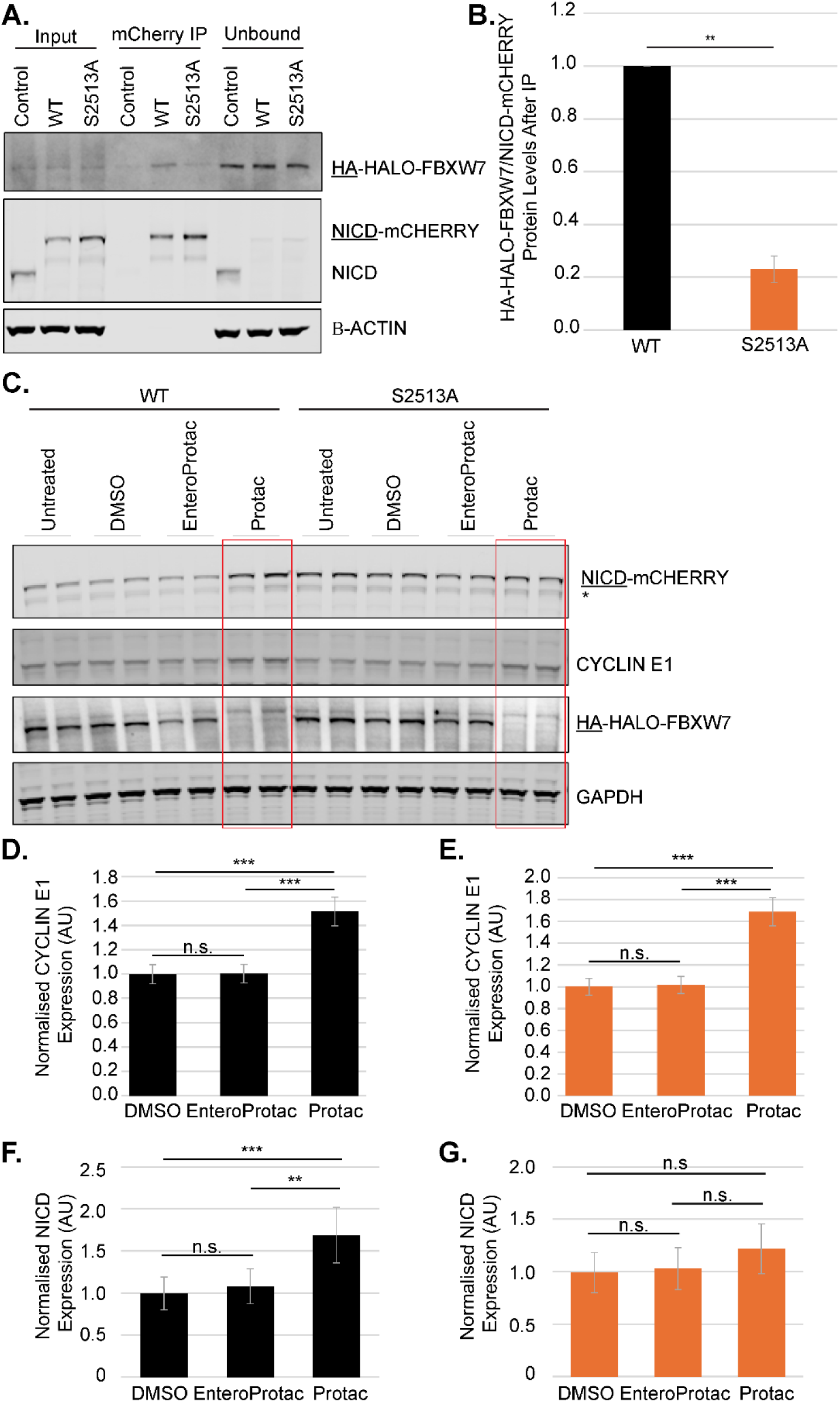
Modification of NOTCH1 S2513 disrupts the interaction of NICD with FBXW7 in the PSM. A) mCHERRY IP on *HA-HALO-FBXW7* (which does not carry an mCHERRY tag and so serves as a negative control), WT and S2513A *NOTCH1* PSM cells. Representative experiment is shown. B) Interacting NICD was quantified (mean +/- s.e.m.) from three biological repeats in A). The amount of interacting HA-HALO-FBXW7 protein for WT NOTCH1 cells was set to 1. S2513A decreases NICD interaction with HA-HALO-FBXW7 by 4.3-fold (t=-14.865, df=2, p=0.0045). C) WT and S2513A *NOTCH1* PSM cells were treated with PROTAC / enteroPROTAC / DMSO for 12 hours. Analysis of expression levels of FBXW7 targets by WB. Representative experiment is shown. D-G) Quantification of four biological repeats (two technical repeats each) of experiment shown in C). The intensity of NICD and CYCLIN E1 bands were normalised to GAPDH and values are displayed relative to vehicle-only DMSO treated values (mean +/- s.e.m.). Depletion of HA-HALO-FBXW7 in WT *NOTCH1* cells significantly increased NICD (Protac v. DMSO t=4.125, df=15, p=0.0009; Protac v. enteroProtac t=3.486, df=15, p=0.0033) and CYCLIN E1 (Protac v. DMSO t=8.15, df=15, p<0.0001; Protac v. enteroProtac t=8.051, df=15, p<0.0001) levels. In S2513A *NOTCH1* PSM cells NICD levels are not responsive to HA-HALO-FBXW7 depletion (Protac v. DMSO t=1.625, df=15, p=0.1249; Protac v. enteroProtac t=1.31, df=15, p=0.2098) but positive control CYCLIN E1 levels are significantly increased (Protac v. DMSO t=10.263,df=15, p<0.0001; Protac v. enteroProtac t=9.901, df=15, p<0.0001). *indicates NICD lacking most of mCHERRY.

### NICD S2513 is required for the control of clock gene mRNA expression in PSM cells

To establish if the S2513A mutation evokes a transcriptional response in PSM cells, the localisation of NICD in PSM cells was assessed using immunofluorescence (IF). This confirmed that the total levels of NICD in S2513A *NOTCH1* PSM cells were increased relative to control cells (Fig 4A). LY411575 treatment served as a control for the NICD immunofluorescence signal and provided additional evidence of the increase of NICD stability in S2513A *NOTCH1* PSM cells compared to WT *NOTCH1* PSM cells. A cell segmentation algorithm (see methods) enabled quantification of the amount of nuclear and cytoplasmic NICD levels. This showed that NICD is predominantly nuclear in both cell lines and that the S2513A mutation caused an increase in nuclear NICD (Fig 4B). To establish whether the increased levels of nuclear NICD in the S2513A *NOTCH1* PSM cells impacts the expression of clock gene mRNAs, WT and S2513A *NOTCH1* iPS cells were differentiated into PSM and clock gene mRNA levels were assayed using qPCR. Significantly increased mRNA levels were detected in the S2513A *NOTCH1* PSM cells for *NRARP*, *HES1* and *LFNG* but not for *HES7* (Fig 4C). The variety in the extent of the response of the different clock genes to increased NICD levels suggests possible mechanistic differences in the regulation of these clock genes, such as differences in the number and orientation of CSL binding sites (Nam et al., 2007). These data demonstrate that the increased levels of NICD caused by the Serine to Alanine mutation in the NICD PEST domain leads to an increase in the expression of NOTCH1 target segmentation clock genes.

**Figure 4.**
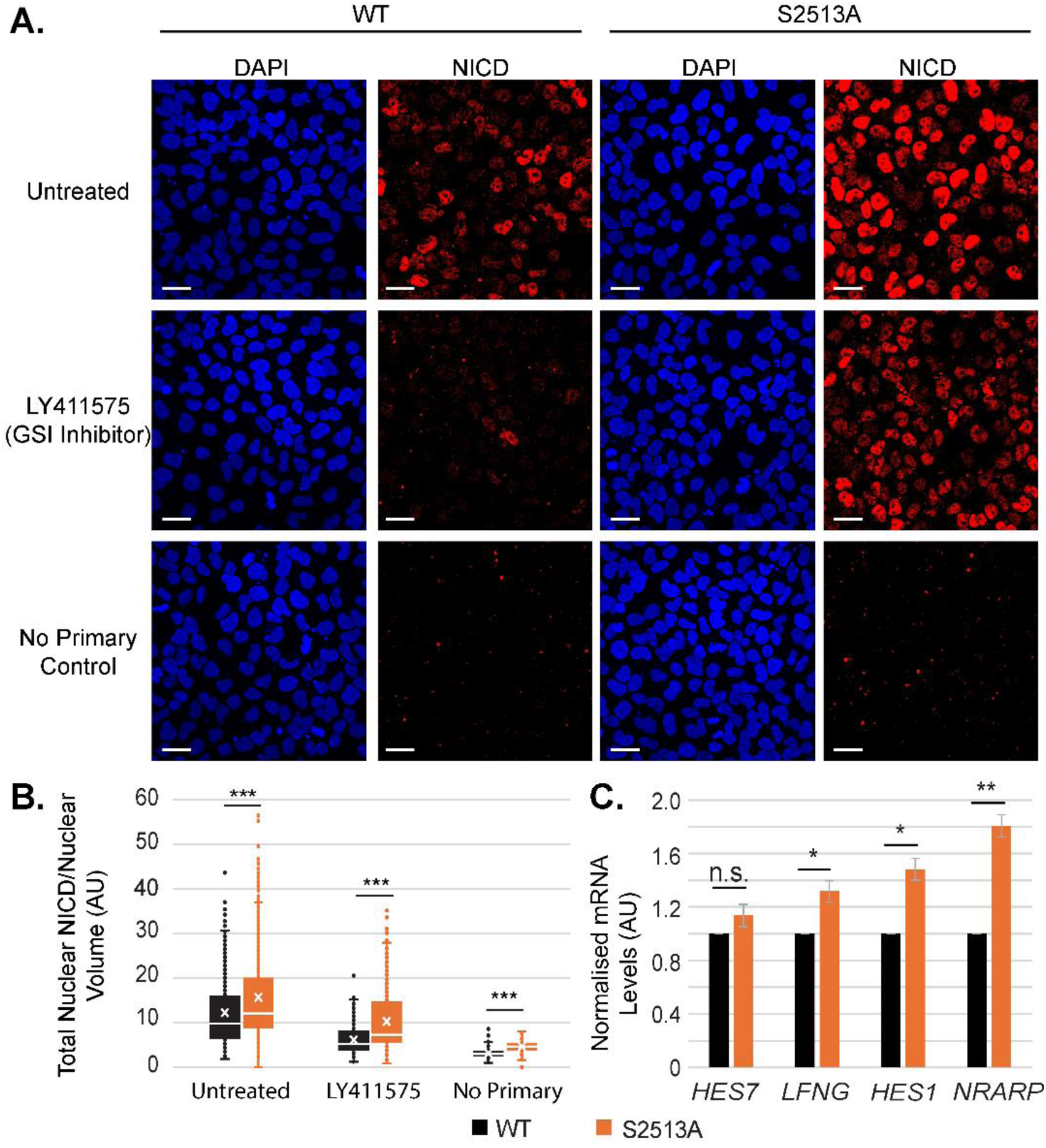
Modification of NOTCH1 S2513 disrupts the regulation of clock gene expression in the PSM. A) WT and S2513A *NOTCH1* IPS cells were differentiated into PSM, selected wells (as indicated) were treated for three hours with GSI inhibitor LY411575, then fixed in 4% PFA/PBS. NICD expression was analysed by IF. Representative experiment shown. Scale bars 25 μm. B) Signal was segmented for nuclear localisation (DAPI positive) and nuclear NICD expression was quantified, corrected for nuclear volume and presented as boxplots from three biological repeats (2-3 FOV per condition). Nuclear NICD levels were increased in both untreated (df=5188, t=10.443, p=<0.0001) and LY treated (df=5188, t=24.41, p=<0.0001) S2513A *NOTCH1* PSM cells when compared with WT NOTCH1 PSM cells. C) WT and S2513A *NOTCH1* iPS cells were differentiated into PSM. RNA was harvested and analysed by qPCR. mRNA levels for four different clock genes were normalised for the expression of a panel of housekeeping genes (*HPRT*, *PPP1CA* and *RNA Pol II*) and analysed for 3 biological repeats (three technical repeats each, mean +/- s.e.m.). mRNA levels of most clock genes tested are significantly increased in S2513A *NOTCH1* PSM cells compared to WT (Hes7 fold change 1.14,df=2, t=1.56, p=0.1291; Lfng fold change 1.32, df=2, t=3.66, p=0.0336; Hes1 fold change 1.48, df=2, t=4.96, p=0.0192; Nrarp fold change 1.81, df=2, t=7.33, p=0.0091).

### NICD S2513 is required for the correct regulation of clock gene oscillations and somitogenesis

To investigate whether the S2513 mutation has a phenotype in PSM tissue, an existing protocol (Sanaki-Matsumiya et al., 2022) was adapted in order to produce somitoids using the WT and S2513A *NOTCH1* hiPSCs (Fig S5A). The key differences with the original protocol are: (i) embryonic bodies were generated prior to differentiation and (ii) retinoic acid was added at the embedding stage, which although not critical for the protocol improved the segmentation (see methods for further details). The WT somitoids exhibited many of the features observed using previous somitoid protocols (Sanaki-Matsumiya et al., 2022; Yamanaka et al., 2023; Miao et al 2023): (i) they formed circular structures that polarised and extended (Fig 5A, 6A, S6A); (ii) they had a well-defined anteroposterior (A-P) axis and distinct PSM region (Fig 5A, 6A, S6A); (iii) they exhibited oscillatory HES7 expression in the PSM region with a period close to that previously reported (link video, Fig 5A, S6A); (iv) they showed propagating waves of expression traversing the P-A axis (link video, Fig 5A/G, S6A). (v) upon embedding in Matrigel they sustained oscillations and produced segments with physical boundaries (link video, Fig 7A, S7). Together, these data indicate that the somitoid system can be used to investigate the effect of S2513A on the segmentation clock and somitoid development. However, it is noted that the formed somite-like structures are heterogenous in shape and size and produce a mixture of single and paired segments, similar to reports using other 3D somite models (Miao and Pourquié, 2024) from which it is challenging to determine segment length, especially since there are changes to the morphology between WT and S2513A somitoids (see below).

**Figure 5.**
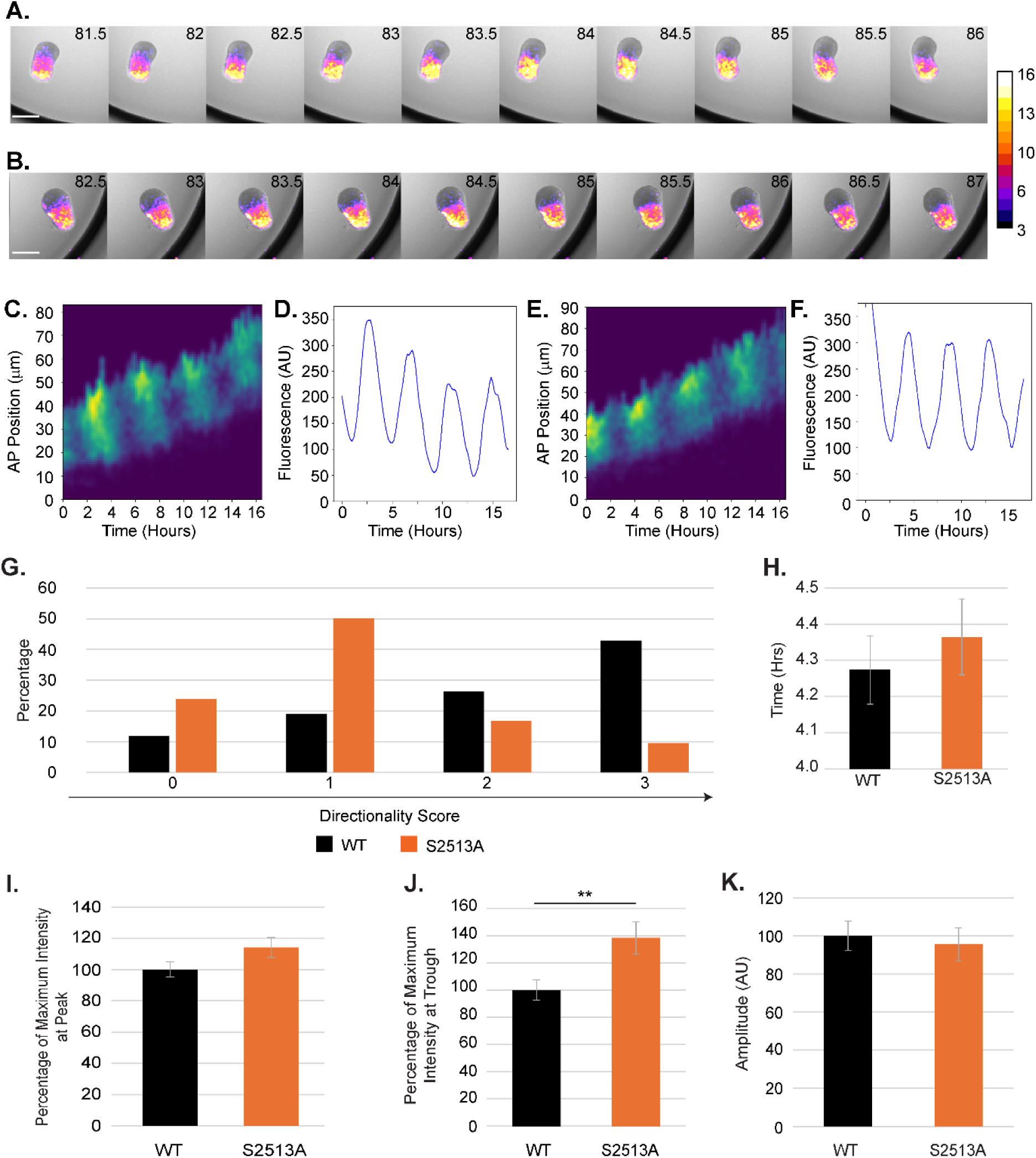
Modification of NOTCH1 S2513 disrupts the segmentation clock in somitoids. A/B) Polarised and elongated WT (A) and S2513A (B) *NOTCH1* somitoids were selected for timelapse imaging. Still images from the timelapse (78-96 hours post differentiation) from representative somitoids are shown. Scale bars 250 μm. Colour bar shows ACHILLES signal intensity. C) Kymograph of the HES7-ACHILLES signal in WT *NOTCH1* somitoids. D) Quantification of the signal intensity of the kymograph in C). E) Kymograph of the HES7-ACHILLES signal in S2513A *NOTCH1* somitoids. F) Quantification of the signal intensity of the kymograph in E). G) Analysis of the kinematics of the oscillations shows that S2513A *NOTCH1* somitoids generally lack P-A directionality. Somitoids from 4 biological repeats (6 each per repeat) were analysed. Directionality score: 0= no/poor oscillations, 1= oscillating – no directional progression, 2= oscillating – some directional progression, 3= oscillating – clear P to A directional progression. H) Quantification of the data obtained from the kymographs shows that the time between signal peaks is not significantly different (increased 6 minutes; t=1, df=33.4, p=0.3244). Graphs show the mean +/- s.e.m. of 4 biological repeats (total of 22 WT and 17 S2513A *NOTCH1* somitoids). I-K) Quantification of the signal intensity from the kymographs, shows an insignificant increase in signal intensity for S2513A NOTCH1 somitoids at the peak 114% (t=1.802, df=34.1, p= 0.0804) and significant increase at the troughs 138% (t=3.004, df=34.1, p= 0.005) of the oscillations. The resulting amplitude has not changed significantly (95%, t=-0.399, df=34.1, p=0.692). Graphs show the mean +/- s.e.m. of 4 biological repeats (total of 22 WT and 17 S2513A *NOTCH1* somitoids).

**Figure 6.**
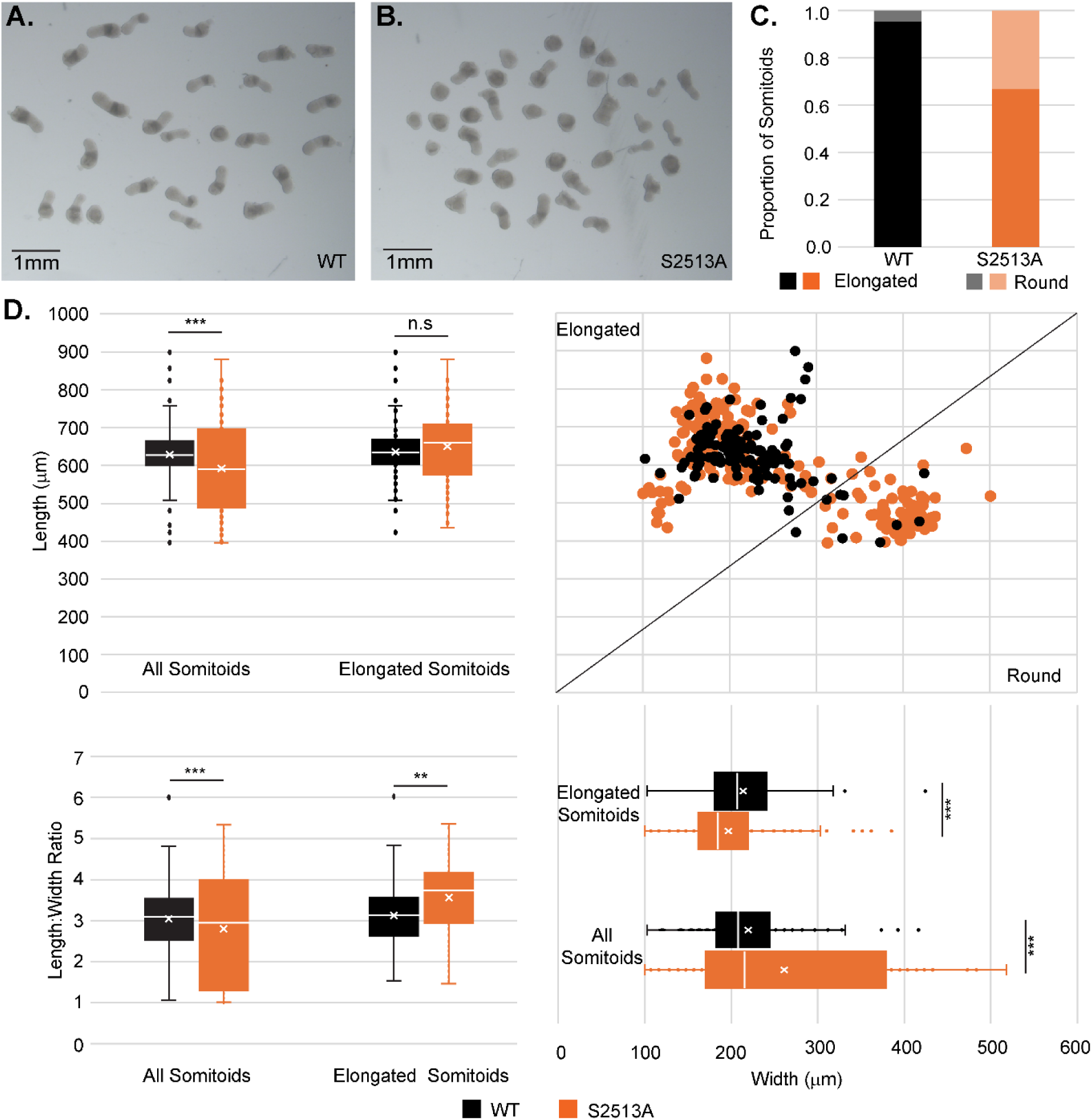
Modification of NOTCH1 S2513 disrupts somitogenesis in somitoids. A/B) Pre-embedding (100 hours post differentiation) WT (A) and S2513A *NOTCH1* (B) somitoids were imaged. A representative experiment is shown. Scale bars 1 mm. C) Somitoid shape was scored. Mean +/- s.e.m. of 7 biological repeats is shown. Almost all WT somitoids elongate successfully whilst 34% of S2513A *NOTCH1* somitoids fail to elongate (Chi-squared=138.65, df=1, p<0.0001). D) Length and width of all somitoids (regardless of shape) for 3 biological repeats was measured confirming the observations made in C) (length: fold change 0.94x, df=320, t=3.367, p=0.0009; width: fold change 1.19x, df=296, t=-4.552, p<0.0001). The calculations were repeated for the somitoids annotated as elongated showing that the elongated WT and S2513A NOTCH1 somitoids have slightly different dimensions, maintaining the same length but being significantly narrower (length: fold change 1.03x, df=238, t=-1.614, p=0.1080; width: fold change 0.92x, df=244, t=2.924, p=0.0038). Line indicates boundary of somitoids classed as elongated and round. The length:width ratio was calculated for both all and elongated somitoids only. The ratio is decreased for all somitoids (fold change 0.92x, df=302, t=3.857, p=0.0001; and increased for elongated somitoids only (fold change 1.14x, df=245, t=-3.614, p=0.0004).

**Figure 7.**
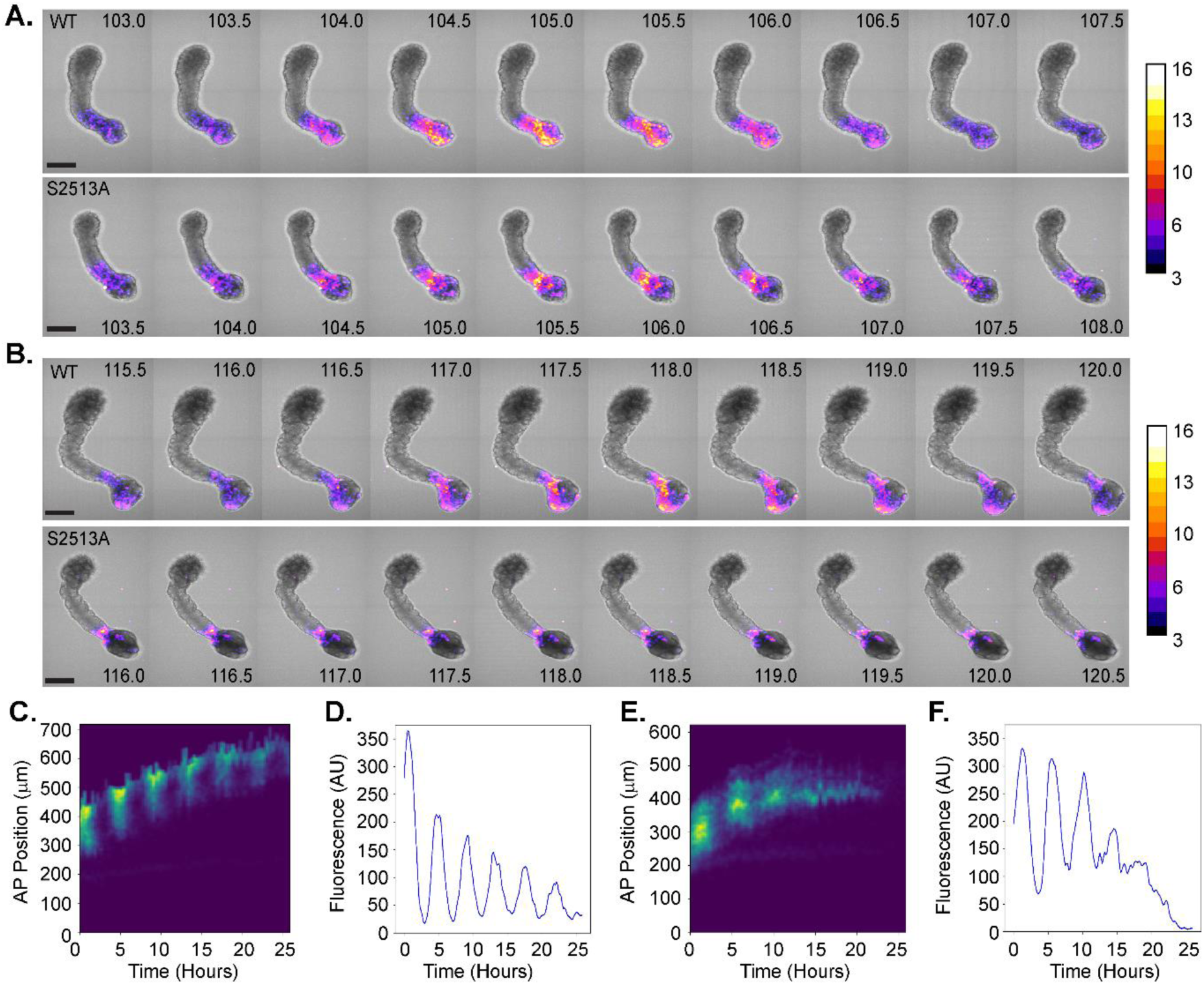
Modification of NOTCH1 S2513 changes the segmentation clock in somitoids. A) Somitoids were embedded ∼97 hours post differentiation and subjected to timelapse imaging (100-126 hours post differentiation). A selection of the time lapse images showing the first peak for WT (top) and S2513A (bottom) *NOTCH1* somitoids. B) A selection of images from the same somitoids as shown in A) showing the fourth peak for WT (top) and S2513A (bottom) *NOTCH1* somitoids. The mutation caused a dampening of the oscillations. Representative selection of images of 4 biological repeats are shown (seven somitoids each per biological repeat). Scale bars 250 μm. Colour bar shows ACHILLES signal intensity. C) Kymograph of WT *NOTCH1* somitoid. D) Quantification of the signal intensity of the kymographs in C). E) Kymograph of S2513A *NOTCH1* somitoid. F) Quantification of the signal intensity of the kymographs in E).

To investigate the effect of S2513A mutation on somitoid development, timelapse imaging (78-96 hours post differentiation) was performed on both WT and S2513A *NOTCH1* somitoids (links videos, Fig 5A-B, S6). Imaging of the Hes7-Achilles reporter demonstrated clear temporal oscillations in both WT and S2513A *NOTCH1* somitoids. After generating kymographs to visualise the HES7-ACHILLES signal intensity along the A-P axis (Fig 5C/E), the signal intensity in the anterior PSM was plotted as a time series (Fig 5D/F). Upon comparison of the WT and S2513A *NOTCH1* somitoids, the pattern of the oscillations differs in several aspects: (i) the Hes7-ACHILLES signal intensity appeared as propagating waves of expression traversing the P-A axis of the WT somitoids (link video, Fig 5A/G, S6A), as reported previously (Sanaki-Matsumiya et al., 2022). However, in the S2513A *NOTCH1* somitoids, the ACHILLES signal was still oscillating, but the directionality of the dynamic signal was disturbed. In the WT *NOTCH1* somitoids the ACHILLES signal started in at the posterior end of the PSM and propagated in an anterior direction whilst for the *NOTCH1* S2513A mutant somitoids the signal started in the centre of the PSM region and then propagated in different directions (link video, Fig 5B/G, S6B); (ii) the average oscillatory period of ACHILLES expression is not significantly different in somitoids prior to embedding in S2513A *NOTCH1* somitoids relative to WT *NOTCH1* somitoids (increased by 6 minutes; Fig 5H), however, this increased after embedding (see below); (iii) whilst the WT and S2513A *NOTCH1* PSM somitoids showed a similar average peak in ACHILLES expression (Fig 5I), the signal troughs are higher in the S2513A *NOTCH1* somitoids (Fig 5J) which is compatible with the increased NICD levels in S2513A *NOTCH1* PSM cells. The resulting amplitude is not significantly different (Fig 5K). Altogether, the timelapse data showed changes to the oscillatory pattern of clock gene expression in the S2513A *NOTCH1* somitoids compared to WT somitoids.

At 100 hours after differentiation (prior to embedding in Matrigel), the WT and the S2513A *NOTCH1* somitoids show different morphological features. The S2513A *NOTCH1* somitoids exhibit reduced polarisation and elongation (67% compared to 95% in WT somitoids) (Fig 6A-C). The S2513A *NOTCH1* somitoids are shorter and wider than the WT *NOTCH1* somitoids (Fig 6D). This analysis does not take the different phenotypes (round versus elongated) into account. When only considering the elongated somitoids, the S2513A *NOTCH1* somitoids are of similar length but thinner than the WT *NOTCH1* somitoids (Fig 6D) resulting in an increased length:width ratio, suggesting that even the elongated S2513A *NOTCH1* somitoids are bearing the effects of disturbed *NOTCH1* signalling. Together, these data show that mutating the NOTCH1 S2513 residue results in a morphological phenotype.

To investigate the role of the local mechanical environment on the phenotype of the S2513A *NOTCH1* somitoids, polarised and elongated somitoids were selected and embedded in Matrigel and analysed by time lapse imaging between 100-126 hours post-differentiation (Fig 7A-B, S7, S8). Oscillation dynamics were analysed in the same way as the unembedded somitoids using kymographs (Fig 7C/E) and oscillation plots (Fig 7D/F). The WT *NOTCH1* somitoids display clear oscillations throughout this timeframe, however the S2513A *NOTCH1* somitoids exhibited damped oscillations and frequently failed to maintain oscillations until the end of the imaging period (Fig 8A, S7, S8). Analysis of the kymographs and oscillation plots showed similar results to somitoids prior to embedding, however, the changes were more pronounced: the time between peaks has increased significantly by 20 min for S2513A *NOTCH1* somitoids (Fig 8B). Even though the signal intensity at the peak of the oscillations is not significantly changed (Fig 8C), the S2513A NOTCH1 somitoids maintain a higher level of HES7-ACHILLES at the troughs of the oscillation cycle (Fig 8D), again indicating that higher NICD levels, due to increased NICD stability, are affecting clock gene oscillations. This resulted in a decrease in the amplitude between peaks and troughs (∼2-fold, Fig 8E). Moreover, even though 88 % of WT *NOTCH1* somitoids form paired somites, S2513A *NOTCH1* somitoids have a reduced ability to form paired somites (24%), generating single somites instead (Fig 8F-G). When performing this experiment without RA the results are similar (Fig S9), however, the morphology of the somitoids is not as well defined as in the presence of Retionic Acid (both WT and S2513A *NOTCH1* somitoids) and there is a higher rate of poorly developed somitoids that do not form clear somites (especially for the *S2513A* NOTCH1 somitoids). Altogether, we have identified qualitative and quantitative differences in HES7-ACHILLES expression dynamics as well as changes in somitoid morphology as a result of the *NOTCH1* S2513 mutation.

**Figure 8.**
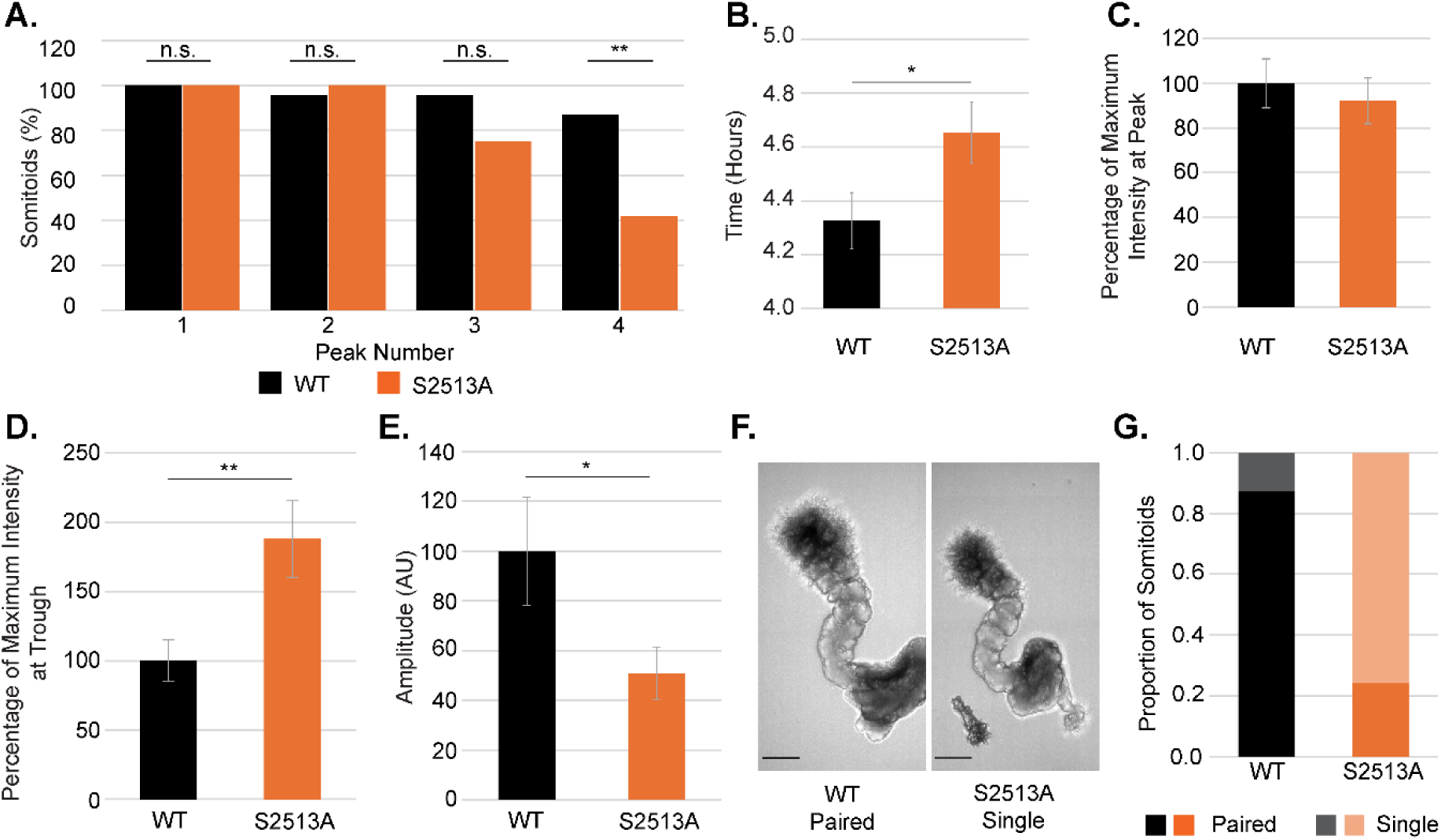
Modification of NOTCH1 S2513 perturbs the segmentation clock in somitoids. A) all somitoids analysed were scored for the number of peaks detected on the oscillation plots. S2513A *NOTCH1* somitoids display fewer peaks than WT *NOTCH1* somitoids with the proportion in the S2513A *NOTCH1* somitoids continuing to 4 peaks significantly smaller than for WT *NOTCH1* somitoids (Chi-squared=8.56, df=1, p=0.0034). Graphs show the mean +/- s.e.m. of 4 biological repeats (total of 23 WT and 24 S2513A *NOTCH1* somitoids). B) Quantification of the data obtained from the kymographs in Fig 7C and 7E demonstrates that the time between signal peaks is increased by 20 minutes in S2513A *NOTCH1* somitoids (t=2.129, df=40.8, p=0.0393). Graphs show the mean +/- s.e.m. of 4 biological repeats for periods following the first three peaks (total of 22 WT and 22 S2513A *NOTCH1* somitoids). C-E) Quantification of the signal intensity from the kymographs, shows an insignificant decrease in signal intensity in S2513A NOTCH1 somitoids at the peak of the oscillations (92%, t=-0.526, df=124, p=0.6001) whilst the signal intensity of the troughs is increased in S2513A *NOTCH1* somitoids compared to WT *NOTCH1* somitoids (188%, t=3.003, df=41.9, p=0.0045). The resulting amplitude between peaks and troughs is decreased in S2513A *NOTCH1* somitoids (51%, t=-2.275, df=40, p=0.0284). Graphs show the mean +/- s.e.m. of 4 biological repeats for the first three peaks and subsequent troughs (total of 22 WT and 22 S2513A *NOTCH1* somitoids). F) Representative images of WT (left) and S2513A (right) NOTCH1 somitoids 126 hours post differentiation. Scale bars 200 μm. G) Analysis of the morphology of WT and S2513A somitoids 126 hours post differentiation. 88% of WT *NOTCH1* somitoids have paired somites, for S2513A *NOTCH1* somitoids this is 24%. 21 WT *NOTCH1* and 23 S2513A *NOTCH1* were scored.

## Discussion

The interaction between FBXW7 and NICD, and subsequent regulation of NICD stability, is critical for several key cellular processes, such as the determination of cell fate, homeostasis, cell differentiation and proliferation (review Kar et al., 2021) and has been shown to be disturbed in several cancer types, e.g. T-ALL, melanoma, breast, salivary gland and colon cancer (review Kar et al., 2021). However, very little was known about the prevalence and/or the significance of the FBXW7-NICD interaction during embryogenesis. Here we have determined that the serine residue at position 2513 in the PEST domain of NOTCH1 regulates NICD stability in PSM cells via the interaction with FBXW7. Moreover, a Serine to Alanine mutation at this residue results in increased stability of NICD and therefore higher NICD and clock gene mRNA levels in PSM cells. This is consistent with observations made in other systems (e.g. Carrieri et al., 2019), however it does not exclude the regulation of NICD via FBXW7 and other E3 ligases via other regions in NICD, as indeed has been shown by others (Broadus et al., 2016).

Furthermore, analysis of the characteristics of the oscillatory patterns the HES7-ACHILLES reporter in hiPSC derived somitoids showed remarkably clear differences between WT and S2513A *NOTCH1* somitoids. The increase in period of clock gene oscillations in the S2513A *NOTCH1* somitoids, is particularly striking considering recent reports showing that protein stability plays an important role in the regulation of timings of several developmental events, such as motor neuron differentiation (Rayon et al., 2021). Indeed, recent reports have shown that species-specific segmentation clock periods are due to differential biochemical reaction speeds (Matsuda et al., 2020). Our data is consistent with these published reports and, furthermore, identifies NICD turnover as one of the key biochemical reactions regulating the clock period.

The WT *NOTCH1* somitoids generated in this study display clock gene oscillations with a period of 4.3 hours which is slightly shorter than previously published (Sanaki-Matsumiya et al., 2022; Miao et al., 2023; Yamanaka et al., 2023). The period for the human segmentation clock reported in the literature does vary dependent on protocol, time window of the time lapse imaging and the spatial location within the PSM selected to analyse the HES7 signal. The somitoids generated using the protocol without the modifications we applied in our study were reported to have a period of 4.5+/-0.2 hours for the anterior PSM when analysed in the same time window as used here and 4.9+/-0.1 hours when that time window was further extended (based on Source Data Sanaki-Matsumiya et al., 2022). Segmentoids (Miao et al., 2023) and axioloids (Yamanaka et al., 2023) generated using different protocols displayed segmentation clock periods reported as 4.6 and approximately 5 hours, respectively. The period reported in this manuscript for WT NOTCH1 somitoids is very similar to the period we observed using HES7-ACHILLES somitoids generated using the same protocol. This suggests that the introduction of the mCHERRY tag is not a contributory factor for the slightly reduced segmentation clock period compared to previously published segmentation clock periods.

Given the number of dynamic unstable components in the feedback loops driving the segmentation clock there are likely to be a variety of mechanisms involved in regulating segmentation clock periodicity. Indeed, a recent study showed that PSM-like cells induced from mouse, rabbit, marmoset, cattle, rhinoceros, or human stem cells display accelerated HES7 oscillations following the removal of the intron in the HES7 gene (Lázaro et al., 2023). The effect of intron removal appears to be proportional to the HES7 oscillatory period of the species in question: the longer the period the bigger the effect of the intron removal (Lázaro et al., 2023). Moreover, inhibition of NOTCH signalling with DAPT was reported to result in a dampening and subsequent loss of HES7 oscillations in human iPS derived PSM cells and somitoids (Diaz-Cuadros at al., 2020; Sanaki-Matsumiya et al., 2022). Dampening of oscillations has also been observed with elevated NICD levels in transgenic zebrafish embryos (Őzbudak and Lewis, 2008) and in DLL1 mouse mutants (Shimojo et al., 2016).

It is nevertheless remarkable that we observed that the directionality, periodicity, intensity and dampening of the HES7-ACHILLES oscillations as well as somite formation were altered due to one single point mutation in NICD. These data suggest that NICD turnover is a central player in the regulation of dynamic NOTCH clock gene mRNA expression in the PSM. Furthermore, as NICD harbours several other potential FBXW7 degrons, it is surprising that, in the context of the S2513A point mutation, there is no effective compensation and rescue of the NICD/FBXW7 interaction through some of the other sites on the NICD PEST domain. This does strongly suggest that, in the context of the PSM, S2513 is a key residue for NICD/FBXW7 interaction and regulation of NICD stability.

In addition to the effect on oscillatory clock gene expression in the PSM, the somitoid data revealed two distinct phenotypes amongst the S2513A *NOTCH1* somitoids: (i) somitoids that polarise, elongate and form somites which closely resemble WT *NOTCH1* somitoids; and (ii) somitoids that fail to elongate/polarise and therefore do not form somites. This suggests the data obtained with 2D culture might also be masking two distinct phenotypes, which is not immediately apparent as all measurements of protein and mRNA levels on the 2D cultured cells are averaged across the whole population of cells. This implies that the impact of the NOTCH1 S2513A mutation on NICD protein and clock target gene mRNA levels might exceed the effects measured here (in cells that give rise to phenotype (ii) above). These observations also demonstrate the power of the somitoid model to analyse the impact of mutations.

Selection of S2513A somitoids at the point of embedding that look similar to the WT somitoids might have inadvertently resulted in an underestimation of the effect of the S2513A mutation. The strongest effects of the S2513A mutation on somitoid development are seen in the percentage of somitoids that elongate and polarise (WT 95%, S2513A 66%), the percentage of somitoids that maintain oscillations until the end of the timelapse imaging after embedding (WT 86%, S2513A 43%) and the percentage of somitoids that generate paired somites (WT 88%, S2513A 24%). These percentages underestimate the full effect of the S2513A mutation on oscillation dynamics and somitoid development as only somitoids that were clearly elongated and polarised were selected for the timelapse experiments. The S2513A somitoids that still maintain oscillations at 126 hours post differentiation as a percentage of all S2513A somitoids is only 28%, compared to 82% for WT somitoids, whilst the generation of paired somites occurs in 84% of WT but only 16% of S2513A *NOTCH1* somitoids.

Both the reduction of NICD and the increase of NICD levels have been shown to disrupt segmentation previously (Őzbudak and Lewis, 2008). Altogether, somitoids with the S2513A mutation can develop, elongate, polarise, display clock gene oscillations and produce paired somites, however, the success rate of achieving this when carrying this mutation is low, and we report statistically significant abnormalities in six distinct aspects of the somitoid phenotype in the S2513A *NOTCH1* somitoids (Fig 9).

**Figure 9.**
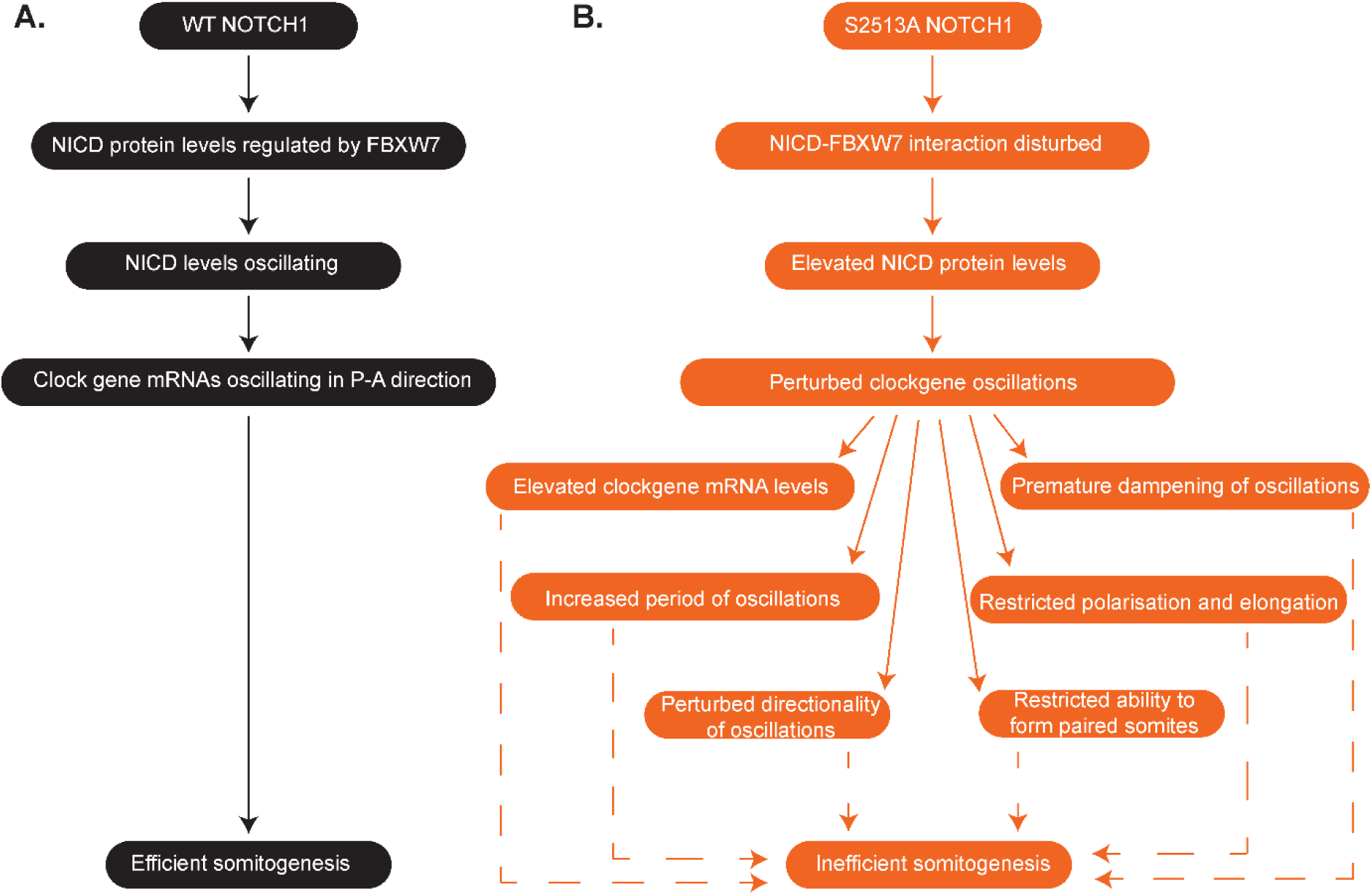
Impact of NOTCH S2513A mutation on somitogenesis. A) In WT *NOTCH1* somitoids NICD levels are regulated by FBXW7, resulting in oscillating NICD protein and clock gene mRNA levels and efficient somitogenesis. B) In S2513A *NOTCH1* somitoids the NICD-FBXW7 interaction is disturbed, resulting in elevated NICD protein levels and perturbed clock gene expression: clock gene mRNA levels are elevated, period of the oscillations is increased, directionality of the kinematic waves is perturbed, oscillations are prematurely dampened, and elongation, polarisation and the formation of paired somites is restricted, together resulting in inefficient somitogenesis.

As discussed, our somitoid protocol generates segments that are heterogenous in shape and size and produce a mixture of single and paired segments, similar to reports using other 3D somite models (Miao and Pourquié, 2024), making somite size measurements between the WT and S2513A mutant cell lines, with 20 min variation in clock gene oscillation period, too variable to draw strong conclusions. Despite this, somitoids generated using human iPS cells provide a powerful model system for the study of human development and disease. Previous studies of the segmentation clock have shown clear differences between species in the regulatory mechanism, e.g. NOTCH is a core clock component in mouse embryos but merely required for synchronisation in zebrafish embryos (reviewed by Yabe and Takada, 2016; Ramesh and Chu, 2024), emphasizing the need for human model systems.

Altogether, the data presented here provides one of the first steps towards understanding the important contribution that post-translational regulation of key factors within the segmentation clock mechanism play in regulating the oscillatory dynamics of oscillatory clock gene expression. The combination of hiPSC-derived 2D and 3D models currently available provide a powerful set of model systems to allow for more detailed studies regarding the mechanisms and gene regulatory networks underlying human development and disease.

## Materials and methods

### Cell lines, hiPSC maintenance, differentiation and inhibitor treatment

Several hiPSC lines have been used throughout this manuscript: Wibj2 iPS cells (female, https://www.hipsci.org); *HES7-ACHILLES* iPS cells that express a self-cleaving T2A peptide followed by the YFP variant ACHILLES at the end of the *HES7* open reading frame (male, Diaz-Cuadros et al., 2020) and several cell lines with additional CRISPR modifications that have been derived from this *HES7-ACHILLES* iPS cell line: *HA-HALO-FBXW7* cells have an HA and an HALO tag added to the *FBXW7* locus, WT *NOTCH1* cells are based on the *HA-HALO-FBXW7* cells and have an mCHERRY tag added to the *NOTCH1* locus and S2513A *NOTCH1* have the mCHERRY tag as well as a point mutation resulting in the change of Serine 2513 into Alanine (Fig S1).

All hiPSC lines were maintained by the Human Pluripotent Stem Cell Facility at the University of Dundee. Briefly, hiPSCs were maintained in TESR medium (Ludwig et al. 2006) supplemented with bFGF (30 ng/ml) and noggin (10 ng/ml) on Cultrex (R&D Systems, 10 ug/cm^2^). Cells were passaged using TrypLE select (Thermo Fischer Scientific) twice a week and seeded at a density of 30,000 – 50,000 cells/cm2 in TESR medium further supplemented with Y27632 (Tocris, 10 μM). Cells were routinely tested to check for mycoplasma contamination using the MycoAlert Detection kit (Lonza), and aerobic bacteria and fungi by inoculation of conditioned cell culture medium into tryptic soy broth (Millipore). No contamination was detected. hiPSC lines were checked by immunofluorescence for the expression of the pluripotency markers OCT4 and NANOG, and pluripotency was determined by *in vitro* differentiation of the cells into ectoderm (*PAX6* expression, Chambers et al. 2009), endoderm (*SOX17* expression, Wang et al. 2017) and mesoderm (*MSGN1* expression, PSM differentiation protocol).

To differentiate iPS cells into PSM, TC plates were coated with Cultrex (R&D Systems, 10 ug/cm^2^) in DMEM/F12 for at least 1 hr at 37°C / 5% CO_2_. Coating mix was aspirated, cells were plated at 15,000/cm^2^ in TESR supplemented with 30 ng/ml bFGF (Peprotech) and 10 μM Y27632 (Tocris) and incubated at 37°C / 5% CO_2_. The next day differentiation was induced by washing the cells twice in PBS and changing the medium to DMEM/F12 (Gibco) supplemented with 1x ITS (Insulin/Transferrin/Selenium, Gibco), 1x MEM NEAA (Non-essential amino acids, Gibco), 1x Glutamax (Gibco), 3.25 μM CHIR99021 (Tocris) and 0.5 μM LDN193189 (Sigma). Cells were fed every 24 hours using the differentiation medium. Experiments were performed after 3 days of differentiation (PSM). Cells were treated with the following inhibitors: 150 nM LY411575 (1/2/3/4 hrs), 1uM MLN4924 (3 hours), 300 nM PROTAC, 300 nM enteroPROTAC (enzymatically inactive PROTAC) or the equivalent volume of DMSO (6 or 12 hrs). For all experiments iPS control cells were plated in parallel and fed with iPS medium every 24 hours and harvested at the same time as the PSM cells. These cells and untreated PSM cells were then analysed by WB to ascertain that the differentiation was effective (data not shown).

### hiPSC derived somitoids

Somitoids were generated using a protocol adapted from the method described by Sanaki-Matsumiya et al., 2022 (Fig S5A). Human iPS cells were seeded into v-bottom plates using 250 cells in 100 μl of TESR medium supplemented with 30 ng/ml bFGF (Peprotech), 10 ng/ml noggin (Peprotech) and 10 μM Y27632 (Tocris) per well. The plate was centrifugated at 300xg for 5 minutes to compact the cells and incubated at 37°C / 5% CO_2_ for ∼47 hrs. The resulting embryonic bodies were transferred to a 60 mm dish and washed twice with DMEM/F12 (Gibco) and subsequently ∼48 embryonic bodies were transferred to a 35 mm dish with 2 ml somitoid induction base medium (SIB: DMEM/F12 (Gibco) supplemented with 1x N2 (Gibco) and 1x B27 (Gibco) supplements, 1x MEM NEAA (Non-essential amino acids, Gibco), 1x Glutamax (Gibco), 1x sodium pyruvate (Gibco), 1x penstrep (Gibco)) supplemented with 10 μM SB431542 (Sigma), 10 μM CHIR99021 (Tocris), 2 μM DMH-1 (Tocris) and 20 ng/ml bFGF (Peprotech)) and incubated at 37°C / 5% CO_2_. Medium was replaced after 24 hours. For the next two feeds at 48 and 72 hours, the somitoids were fed with SIB. Approximately 96 hours after differentiation individual somitoids were embedded in 10% Matrigel in SIB supplemented with 150 nM Retinoic Acid (Sigma) in 15 well ibidi slides coated with 100% Matrigel / 150 nM Retinoic Acid (Sigma) to allow for segment formation. Exception to this protocol are the somitoids in Fig S9: they were grown in the absence of Retinoic Acid and either on 50% or 100% Matrigel coated slides.

### CRISPR modification of hiPSC

Snapgene maps for *FBXW7* and *NOTCH1* loci were compiled by directly importing NCBI contig data; Chromosome 4 into Snapgene; NC_000004.12 (152320544..152536092, complement) and Chromosome 9, NC_000009.12 (136494433..136546048, complement) respectively. Ensembl was used to cross-reference and confirm the annotation of all known transcript variants. Prospective guides with low off-targeting scores and situated close to the ATG start codon of *FBXW7* and S2513 residue of *NOTCH1* were identified using a Sanger webtool (https://wge.stemcell.sanger.ac.uk/find_crisprs).

For *HA-HALO-FBXW7* donor construction WT Cas9 and a single guide (G2 5’-AGCAAAAGACGACGAACTGG) were used. For both WT and S2513A *NOTCH1* nickase Cas9 and a guide pair (left guide 5’-GTCAGGGGACTCAGGGGACG, right guide 5’-CTCAGGGGACGGGGTGAGGA) were used. Complementary oligos with BbsI compatible overhangs were designed according the Zhang method (Cong et al., 2013), annealed and the resulting dsDNA guide inserts ligated into BbsI-digested target vectors; nickase left guides were cloned into the spCas9 D10A expressing vector pX335 (Addgene #42335), right guides into the puromycin selectable plasmid pBABED P U6 (DU48788, MRC-PPU Reagents and Services, University of Dundee) and unpaired guides cloned into the spCas9-puro vector pX459 (Addgene #62988) yielding plasmids DU69619, DU69620, DU69621, DU69622, DU57766, DU57768, DU57767 and DU57784 respectively. A *pMA PURO-2A-HA-HALO-FBXW7* donor (DU69658) comprising 500 bp homology arms flanking the transgene insert and containing sufficient silent nucleotide changes to prevent guide recognition was synthesised by GeneArt.

WT and S2513A *NOTCH1* donor construction: The GFP markers in existing donors DU64733 (*pMK-RQ NOTCH1* Cter S2513A 6Gly GFP) and DU64770 (*pMK-RQ NOTCH1* wt Cter 6Gly GFP) were swapped for mCHERRY via Gibson assembly: mCHERRY insert PCR amplified from DU74115 and *NOTCH1* vector backbones amplified from DU64733 and DU64770. For primers see Table S3. PCRs were performed in 50 μl reactions using KOD Hot Start according to manufacturers’ recommendations, 1 μl of DpnI was added to each PCR to degrade template DNAs and, following incubation for 30 minutes at 37°C and 15 minutes at 65°C, the products were cleaned using a PCR cleanup kit (QIAgen) and eluted in 50 μl H_2_O. Gibson assemblies were performed, combining 100 ng of cleaned mCHERRY insert with 100 ng of the appropriate donor backbone. Following incubation for 1 hour at 50°C, 2 μl of each Gibson reaction was then transformed into chemically competent DH5α cells (MRC-PPU Reagents and Services, University of Dundee).

Electroporations of iPS cells were performed using a Neon electroporator (Thermo Fischer Scientific) using 10 μl tips. Briefly, cells were passaged using TrypLE select and resuspended in TESR medium supplemented with Y27632 (10 μM). 1×10^6^ cells were collected by centrifugation at 300xg for 2 minutes, washed with PBS, collected by centrifugation at 300xg for 2 minutes, then resuspended in 10 μl electroporation buffer R. 1 μg of plasmid repair template and 1 μg of guide plasmids (0.5 μg each for the nickase pair for NOTCH1) were then added in a volume of 2 μl. Cells were then electroporated at 1150v, 30 mSec, 2 pulses then plated on Geltrex (Corning) coated dishes (10 μg/cm2) in TESR medium supplemented with Y27632.

For HA-HALO-*FBXW7* clones: after 24 hours the medium was replaced with fresh TESR medium and cells grown until confluent (medium replaced every 24 hours). Cells were then dispersed, and 400 cells seeded into 60 mm dishes to generate monoclonal cell lines. When colonies were ∼2-3mm in diameter they were picked using 3.2 mm cloning discs (Bel Art Laboratories) soaked in TrypLE select and seeded into 96 well plates. Edited clones were identified by staining the cells with HALO tag TMRDirect ligand (Promega) according to the manufacturer’s recommendations, screened (see below), expanded, banked and frozen.

For WT and S2513A *NOTCH1* clones: after 24 hours the medium was replaced with fresh TESR medium and cells grown until confluent (medium replaced every 24 hours). mCHERRY positive cells were purified by Fluorescence Activated Cell Sorting (FACS) as a bulk population. FACS was performed using an MA900 cell sorter from Sony Biotechnology, equipped with a 130 µm nozzle. Forward angle light scatter (FSC) and back scatter (BSC) were generated using a 488nm laser and detected using 488±17nm band pass filters. Cells were distinguished from debris based on FSC-Area(A) and SSC-A measurements. Single cells were distinguished from doublets and clumps based on FSC-A and FSC-Width (W) measurements. mCHERRY was excited by a 561nm laser and fluorescence detected using 617±30 band pass filter. Positive cells were identified by assessing the background autofluorescence of control (untransfected) cells which did not express mCHERRY. Cells were collected into TESR medium containing bFGF (30 ng/ml), noggin (10 ng/ml) and Y27632 (10 µM) using the MA900 cell sorter in semi-purity mode and seeded back into a 6 well. Individual clones were then picked using 3.2 mm cloning discs (Bel Art Laboratories) soaked in TrypLE select when they were ∼2-3mm in diameter and seeded into 96 well plates. After screening (see below) selected clones were expanded, banked and frozen.

### Screening CRISPR modified clones

For CRISPR modified hiPSCs gDNA was purified using the GenElute Mammalian Genomic DNA Miniprep Kit (Sigma) following the manufacturer’s protocol. The insert and surrounding region were amplified by PCR using KOD Hot Start DNA Polymerase (Novagen) according to the manufacturer’s protocol with the following modifications: DMSO was added to a final concentration of 6% (v/v) and gDNA at 4 ng/μl. Sequencing was performed by MRC-PPU Reagents and Services (University of Dundee). Both strands were sequenced to confirm correct insertion of modifications/tags (data not shown). Primers used for PCR and sequencing are listed in Table S3. Cell line authenticity was confirmed by the European Collection of Cell Cultures (ECACC) using their AuthentiCell testing kit for all new cell lines. They were screened for any contamination as described in the section on hiPSC maintenance. No contamination was detected.

### RNA purification and qPCR analysis of clock gene expression

WT and S2513A *NOTCH1* iPS cells were differentiated into PSM. RNA was harvested from 3 separate wells for each cell line per experiment using RNeasy mini columns (QIAGEN) combined with QIAshredder columns (QIAGEN) to homogenise the samples. RNA purification was according to the manufacturer’s protocol with the addition of a DNAseI incubation for 15 min at RT using 27 KU of DNaseI in the provided RDD buffer (QIAGEN). The eluted RNA was quantified and analysed on a Nanodrop (Thermo Fisher Scientific). A260/280 values for all samples were between 2.0 and 2.1. The integrity of the RNA was checked by running it on a 2% agarose in TBE gel and clear 28S and 18S bands were observed for all samples without any visible signs of degradation of the RNA. cDNA was generated by denaturing 500 ng of RNA with 150 ng random hexamers (Thermo Fisher Scientific) and 10 pmol dNTP mix (Thermo Fisher Scientific) for 5 min at 65°C and snap cooling on ice. The RNA was reverse transcribed in 1^st^ strand buffer supplemented with 5mM DTT using 200 U Superscript III (Thermo Fisher Scientific) in the presence of 20 U SuperaseIn (Thermo Fisher Scientific) by incubating for 5 min at 25°C, 60 min at 50°C, 15 min at 70°C before cooling to 4°C. The resulting cDNA was used for qPCR with 1x Luna Universal qPCR Mastermix (NEB) and 200 nM of forward and reversed primers using a BioRad CFXConnect qPCR machine with CFX Maestro software and the following program: 20 sec 95°C, 40 cycles of 3 sec at 95°C and 30sec at 60°C, followed by a melt curve. The melt curves for all primer pairs showed one peak only and when analysed on 1% agarose gel in TAE showed a single band only. qPCRs were performed with primers for the clock genes *HES7*, *LNFG*, *HES1* and *NRARP* as well as the housekeeping genes *HPRT*, *PPP1CA* and *RNA Pol II* (for primer sequences see Table S3). Data from triplicate wells were analysed using the ddCq method with the results for the clock genes normalised for the average of the housekeeping genes. Cq values for triplicate wells were within 0.5 Cq. Levels of housekeeping mRNAs varied by a maximum of 15% between samples. For all primers the primer efficiency was checked and found to be within 90-110%. No RT and water controls were performed for all experiments and contributed to less than 1% of the relative amounts obtained with the plus RT samples.

### Pluripotency tests

For every new cell line qPCR analysis was performed to establish whether the cells expressed iPS markers (*OCT4* and *NANOG*) and can be differentiated into the three germ layers (ectoderm – *PAX6*, mesoderm – *MSGN1*, endoderm – *SOX17*). To generate cDNA one μg total RNA was reverse transcribed using a high-capacity reverse transcription kit (Thermo Fisher Scientific Cat. no. 4368814) according to the manufacturer’s instructions without RNAse inhibitor. qPCR was performed as for the analysis of clock gene expression. Data was presented as average Cq values from triplicate wells.

### Immunoprecipitation

Beads (HA-HIS Frankenbody or mCHERRY2-HIS, MRC-PPU Reagents and Services, University of Dundee) were resuspended on rollers at 4°C for a minimum of 30 minutes. 50 µl of bead slurry per condition were washed twice in IP buffer (50mM Tris-HCl pH 7.5, 50mM NaCl, 0.5% Triton X-100) before being blocked at 4°C in 2% BSA in IP buffer for 3-4 hours before a final wash step. Meanwhile, cells were treated with 1uM MLN4924 (MRC reagents and Services, University of Dundee) for 3 hours. After treatment cells were washed with PBS and scraped into IP buffer with inhibitors (cOmplete Mini EDTA-free protease inhibitor cocktail (Sigma), 50mM NaF, 1mM Na_3_VO_4_, 1μg/ml microcystin). Lysates were sonicated on high for 6 rounds of 30s on and 30s off (Bioruptor Plus, Diagenode) and incubated on ice for 10 minutes before being centrifuged at 4°C, 20,600 x g for 15 minutes. Supernatant was transferred to a new tube, protein content was quantified via Bradford Assay (Bio-Rad) and 400µg of protein was made up to 200µl in IP buffer plus inhibitors and was added to the blocked/washed beads and incubated at 4°C on rollers for 2 hours. Tubes were centrifuged at 4°C, 425 x g for 1 minute and supernatant collected as unbound. Beads were washed 3 times in IP buffer and finally resuspended in 15µl of 4x SDS-PAGE loading buffer + BME (bromophenol blue, 4% SDS, 250mM Tris-HCl pH 6.8, 40% glycerol, 5% β-mercaptoethanol).

### Cell lysis, protein quantification and western blotting

Cells were lysed in 1.25x SDS-PAGE loading buffer (1.25% SDS, 78mM Tris-HCl pH 6.8, 12.5% glycerol). Lysates were sonicated on high for 12 rounds of 30s on and 30s off (Bioruptor Plus, Diagenode) and incubated on ice for 10 minutes before being centrifuged at 4°C, 20,600 x g for 15 minutes. Supernatant was transferred to a new tube and proteins were quantified by BCA (Thermo Fisher Scientific). Samples were diluted with 4x SDS-PAGE loading buffer + BME to produce equal concentrations for gel loading. Proteins were separated on precast 4-12% NuPAGE gels (Thermo Fisher Scientific) and transferred to 0.45 uM Protran Nitrocellulose (Amersham). Membranes were rinsed in TBST, blocked for at least one hour in 5% milk in TBST, rinsed in TBST and incubated in primary antibody (see Table S4) O/N at 4°C on a rocker. The next day the membranes were washed 3x 5 min in TBST and incubated for at least 2 hours in secondary antibody (see Table S5) at RT on a rocker. Membranes were washed 5x 5 min in TBST prior to imaging on an Odyssey Imaging System (LI-COR) or on a Chemidoc (Biorad, HA and FBXW7 blots only). Bands were quantified using Image Studio software (LI-COR).

### Immunofluorescence (IF) on cells

Monolayer cells were fixed with 4% PFA/PBS at RT for 10 minutes, washed with PBS, permeabilised with 0.1% Triton for 10 minutes, washed with PBS and endogenous peroxidase activity was quenched with 3% H_2_O_2_ solution for 10 minutes. Cells were washed with PBS, blocked using 10% goat serum (Thermo Fisher Scientific) for 1 hour, before incubation with primary antibodies for 1 hour (see Table S4). Following this, cells were washed with PBS before incubation in secondary antibodies (see Table S5) and 1μg/ml DAPI (Sigma) for 1 hour. For NICD IF the signal was boosted with AF594 TSA reaction mix (Invitrogen) made up according to the manufacturer’s instructions and cells were incubated in reaction mix for 10 minutes, followed by incubation in stop reagent (Invitrogen) for 10 minutes. Cells were washed with PBS and stored at 4°C until imaging. All incubation steps were carried out on a rocker at RT.

### Imaging fixed somitoids and cells

Embedded and fixed somitoids were imaged on 15 well ibidi µ-slides using a 10x dry immersion lens. Fixed cells were imaged on 8 well ibidi µ-slides using a 63x oil immersion lens (for NICD) or a 20x dry lens (for NANOG and OCT4). All were imaged on a Leica SP8 Stellaris microscope maintained by the University of Dundee’s Imaging Facility. After allowing the microscope to warm up, up to 3 regions of interest per condition were selected where cells were organised in a monolayer. Selection was carried out using the DAPI filter to prevent NICD signal-related bias. To prevent overexposure, microscope settings for DAPI, ACTB and NICD signals were optimised using “S2513A untreated” conditions which yielded the highest NICD signal. Identically sized z-stacks were imaged with settings kept constant across all imaged conditions.

The quantitative data from the confocal images was generated using in-house Python pipelines. This Confocal data set contains 3D z-stack IF images of three channels: DAPI, ACTB and NICD. The dataset contains two cell types (WT and S2513 *NOTCH1*), and two conditions (untreated and LY411575 treated). The deep learning algorithm, cellpose (Stringer et al., 2021), was used on the DAPI and the ACTB channel to segment the nuclei and the cell outline with the parameters listed in Table S6. This generates marks or regions of interest (ROI) of segmented nuclei and cells. ROIs were only considered if both nucleus and cell outline were segmented. Next, the NICD pixel intensity was measured within each ROI by taking the mean value using *np.mean()* of the pixel intensity. The results were saved as a CSV, with each cell representing a row with cell number and NICD pixel intensity. The CSV output was then analysed using Panda’s library for data formatting and pre-processing, matplotlib, and Seaborn for data visualisation. We acknowledge using GitHub Copilot and ChatGPT to help generate the Python pipelines. The tool was used to help debug, optimise, and provide help with code snippets. The writing code snippets included helping write functions for pre-processing the images and methods to help generate CSV files. The code was then modified and tested to ensure the snippets functioned as intended.

### Time lapse imaging (pre/post embedding)

For the timelapse imaging pre-embedding somitoids that were observed to have begun polarisation were selected approximately 77 hours after differentiation and transferred to a 15 well ibidi µ-slide. For timelapse imaging post-embedding somitoids that were clearly elongated were selected 97 hours after differentiation and transferred to a Matrigel coated 15 well ibidi µ-slide. Slides were imaged using a 10x dry lens on a Leica SP8 Stellaris microscope with an atmospheric control attachment set to 37°C, 5% CO_2_ and left to settle for at least 30 minutes. Measuring EYFP (ACHILLES) and brightfield, 245μm z-stacks for each somitoid were taken at 10-minute intervals over the course of 18/26 hours. The P-A directionality of the ACHILLES oscillations was scored by 3 independent researchers and median scores were used to establish the ability of the somitoids to generate oscillations that travel in a P-A direction within the PSM. Morphology of the somitoids was scored in the same way and averaged scores were plotted to establish the ability of the somitoids to form paired or single somites. Live imaging of the somitoids yielded brightfield and HES7-ACHILLES 4D image stacks (t,z,x,y). The brightfield signal was z-projected and each xy plane was normalised. The cellpose ‘cyto’ algorithm was retrained to segment somitoids using manually annotated images. The centroid and major axis of the segmented somitoid were identified (*skimage-regionprops*). A small sample of points along the major axis were used as seeds for a K-means clustering algorithm (*sklearn-KMeans*) that was used to cluster points on the segmented somitoids. A travelling salesman algorithm was then used to identify the shortest route between the identified K-means clusters. Piecewise linear segments (scipy-interpolate) were used to interpolate the cluster centres. Hence an AP axis was defined. The HES7-ACHILLES fluoresence intensity image was despeckled using a median filter in each xy plane (*ndimage-median filter*). A Gaussian filter was then applied in each xyz plane (*ndimage-gaussian_filter*). The filtered HES7-ACHILLES signal was z-projected. At each point on the identified A-P axis, the signal was averaged along a local direction perpendicular to the AP axis. Hence a kymograph was obtained. A Gaussian filter was applied to the kymograph (*ndimage-gaussian_filter*). The anterior region of the kymograph was defined to be a fixed distance from the posterior end of the AP axis. A time series was generated signal and peaks and troughs were identified (*scipy-signal-find_peaks*). The time elapsed between successive peaks was used to estimate signal period. Image analysis parameters: median filtering neighbourhood (3), gaussian filtering σ (1), Num K means clusters (5), gaussian filtering σ kymograph (1), minimum period (3h) (Fig S5B).

### Analysis somitoids: shape and size

Pre-embedding somitoids were imaged in 35 mm dishes using an AmScope camera attached to a Leica MZ16 microscope. Somitoids were divided into groups based on their shape (elongated or round). Any somitoids that were fused or had resulted from fragmented embryonic bodies were discarded. Images of somitoids were analysed in Fiji using the ‘points’ function to measure the length and the width of the somitoids by calculating the Euclidean distance.

### Statistics

All statistical analyses of experiments on PSM cells have been performed on three or four biological repeats. In most cases the data was treated as a block design with subsampling to correctly account for the technical repeats. A linear mixed effects model was fitted to the logarithm of the data and non-significant factors sequentially excluded. The factorial categories were treated as fixed effects with the technical replication included as a random effect (Fig 1, 2, 3D-G and S4). All models were checked with a variance inflation test to ensure no unconsidered interactions had been excluded from the model. IF image quantifications were analysed using a linear model with nuclear amount corrected for nuclear volume as the reporting variable. Interactions were included for cell type and treatment, but not for the individual batch as this proved to not be significant in model optimisation (Fig 4B). For the qPCR analysis, each biological replicate was treated as an independent measure containing three technical replicates each of which was measured three times. Delta Cq values were corrected using the pool of housekeeping genes as a reference. Variances from the technical repeats were propagated to allow calculation of the t-statistic for the mean delta Cq and p-value determination using only the degrees of freedom from the biological repeats to obtain a conservative estimate of significance (Fig 4C). For the somitoid experiments several biological repeats were performed as indicated in the figure legends. As both wild type and mutant somitoids were treated together in each batch, the raw fluorescence intensities could be used as a reporter value and directly compared. The batch and cell type were therefore treated as fixed effects. The logarithm of the intensity was the reporter value allowing for a normal error model. The individual somitoid was treated as a random effect to control for repeated measures at the different time points (Fig 5, 7 and 8). T-tests were used for the analysis of the IP data and the somitoid size measurements (Fig 3B and 6D-F), Chi-squared tests were used in the analysis of proportions (Fig 6C and 8A). Statistical analysis was performed in R 4.4.1 with the lmer and Emmeans packages.

## Supporting information

movie WT pre embedding somitoid

movie S2513A pre embedding somitoid

movie WT embedded somitoid

movie S2513A embedded somitoid

## Acknowledgements

We would like to thank Olivier Pourquié (Harvard Medical School, Boston, USA) for the kind gift of the *HES7*-ACHILLES cell line. We would like to thank Alessio Ciulli (Centre for Targeted Protein Degradation, University of Dundee) for the kind gift of the HALO-PROTAC and HALO-enteroPROTAC. We would also like to thank Emily Shak and Kim Peters (lab of Kim Dale, University of Dundee) for their contribution to clone screening and sequencing, Febe Ferro (lab of Kim Dale, University of Dundee) and Olga Suska (lab of Vicky Cowling, University of Dundee) for their contribution to primer design as well as the Imaging and FACS facilities at the University of Dundee for their expertise and MRC-PPU for reagents and services.

## Author contributions

H.A.M., J.K.D., P.M., P.D. and S.J. conceptualised the research; L.D., H.A.M. and A.H. optimised the somitoid protocol with additional recommendations from T.E.J.C.N. and K.F.S.; L.D., H.A.M., M.K. and T.J.M and maintained the iPS cells and designed, generated and screened the CRISPR clones and generated the somitoids; H.A.M., A.H., S.J., I.N.H. and J.M.R. performed the experiments; A.H., P.M. and R.L.G. performed imaging and image analysis; D.M.A.M. provided advice and assistance for the statistical analysis; H.A.M. wrote the original draft of the manuscript and all other authors made contributions to the methods section and*/*or provided feedback on the manuscript.

## Supplementary information

**Figure S1 Schematic representation of *HES7-ACHILLES*, *HA-HALO-FBXW7*, WT and S2513A *NOTCH1* cell lines**

*HES7-ACHILLES* cells express the fluorescent protein ACHILLES under control of the *HES7* promoter. For the *HA-HALO-FBXW7* cell line HA and HALO tags were added to the endogenous *FBXW7* locus to enable efficient detection and degradation of FBXW7. To the resulting cell line an mCHERRY tag was added to the endogenous *NOTCH1* locus and *NOTCH1* Serine 2513 was mutated into Alanine.

**Figure S2 iPS marker checks for *HES7-ACHILLES*, *HA-HALO-FBXW7*, WT and S2513A *NOTCH1* cell lines**

iPS cells were analysed by IF for pluripotency markers. All cell lines *HES7-ACHILLES* (A/B); *HA-HALO-FBXW7* (C/D); WT *NOTCH1* (E/F); S2513A *NOTCH1* (G/H) express both markers tested NANOG (A/C/E/G); OCT4 (B/D/F/H). One biological repeat, three fields of view (FOV) each. Representative image shown. Scale bars 25 μm.

**Figure S3 Differentiation checks for *HES7-ACHILLES*, *HA-HALO-FBXW7*, WT and S2513A *NOTCH1* cell lines**

A/B) *HES7-ACHILLES*, *HA-HALO-FBXW7* (A) WT and S2513A *NOTCH1* (B) iPS cells were differentiated into PSM cells. Every 24 hours samples were collected for WB analysis. iPS and PSM marker expression is very similar for all cell lines. NICD WB shows two bands for WT and S2513A NOTCH1 PSM cells: top band representing NICD-linker-mCHERRY; bottom band representing NICD-linker-small part of mCHERRY. Representative experiment shown of two biological repeats. C) WT and S2513A *NOTCH1 iPS* cells were differentiated into PSM or neuroectoderm (NE) cells. The S2513A mutation prevented efficient differentiation into NE but not PSM cells. Representative experiment shown of three biological repeats. *indicates NICD lacking most of mCHERRY.

**Figure S4 FBXW7 target expression and PROTAC effectiveness test for HA-HALO-FBXW7 cells**

A) *HES7-ACHILLES* and *HA-HALO-FBXW7* cells were differentiated into PSM cells. Expression levels of proteins targeted by FBXW7 for degradation were analysed by WB. Representative experiment is shown. B) Four biological repeats (three technical repeats for each) of A) were quantified (mean +/- s.e.m.). NICD and CYCLIN E1 levels were normalised to GAPDH. No effect of the introduction of the *HA-HALO* tag to the *FBXW7* locus was observed for NICD (fold change 0.97x, t=0.425, df=3, p=0.6996) or CYCLIN E1 (fold change 0.98x, t=0.739, df=3, p=0.5135) levels. C) *HA-HALO-FBXW7* iPS cells were treated with PROTAC / enteroPROTAC / DMSO for six hours. Cell lysates were subjected to HA-IP and resulting samples were analysed by WB. PROTAC treatment efficiently depletes FBXW7 from the cells. The FBXW7 antibody can detect HA-HALO-FBXW7 after IP but not in total lysate. Representative experiment of three biological repeats is shown.

**Figure S5 generation of somitoids and timelapse imaging analysis**

A) Timeline somitoid protocol. SIB = somitoid induction media. B) Flowchart depicting the steps required for the timelapse image analysis.

**Figure S6 Still images from the timelapse imaging (78-96 hours post differentiation)**

A) WT NOTCH1 somitoids. B) S2513A NOTCH1 somitoids. Representative somitoids are shown. Scale bars 250 μm. Colour bar shows ACHILLES signal intensity. Time stamp reflects time in hours from differentiation.

**Figure S7 Still images from the timelapse imaging (100-126 hours post differentiation) – WT NOTCH1 somitoids**

Representative somitoids are shown. Scale bars 250 μm. Colour bar shows ACHILLES signal intensity. Time stamp reflects time in hours from differentiation.

**Figure S8 Still images from the timelapse imaging (100-126 hours post differentiation) – S2513A NOTCH1 somitoids**

Representative somitoids are shown. Scale bars 250 μm. Colour bar shows ACHILLES signal intensity. Time stamp reflects time in hours from differentiation.

**Figure S9 S2513A NOTCH1 somitoids have a reduced ability to form paired somites**

Analysis of the morphology of WT and S2513A somitoids 120 hours post differentiation in the absence of RA. 63% of WT *NOTCH1* somitoids have paired somites, for S2513A *NOTCH1* somitoids this is 25%. 84 WT NOTCH1 and 88 S2513A NOTCH1 were scored.

## Supplementary Tables

Table S1 Expression of iPS markers

Table S2 Expression of differentiation markers

Table S3 Primers

Table S4 Primary antibodies

Table S5 Secondary antibodies

Table S6 Cellpose parameters

