## Supplementary figures and images for "NOTCH1 S2513 is critical for the regulation of NICD levels impacting the segmentation clock in hiPSC-derived PSM cells and somitoids"

### Supplemental figures

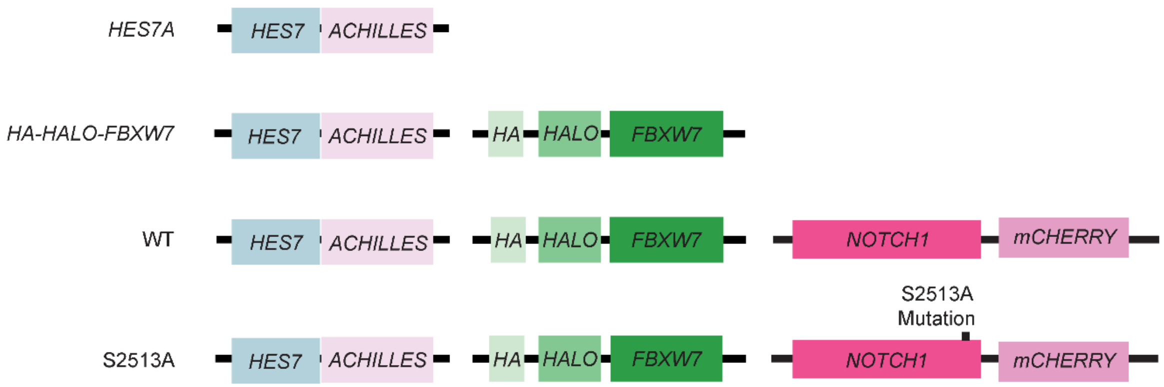

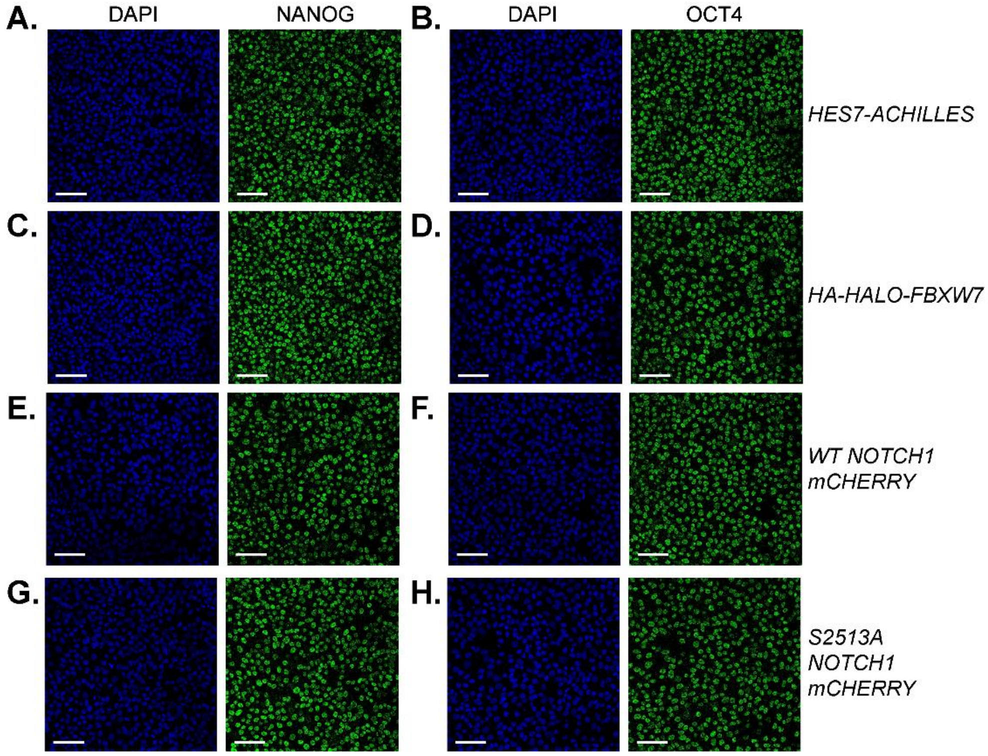

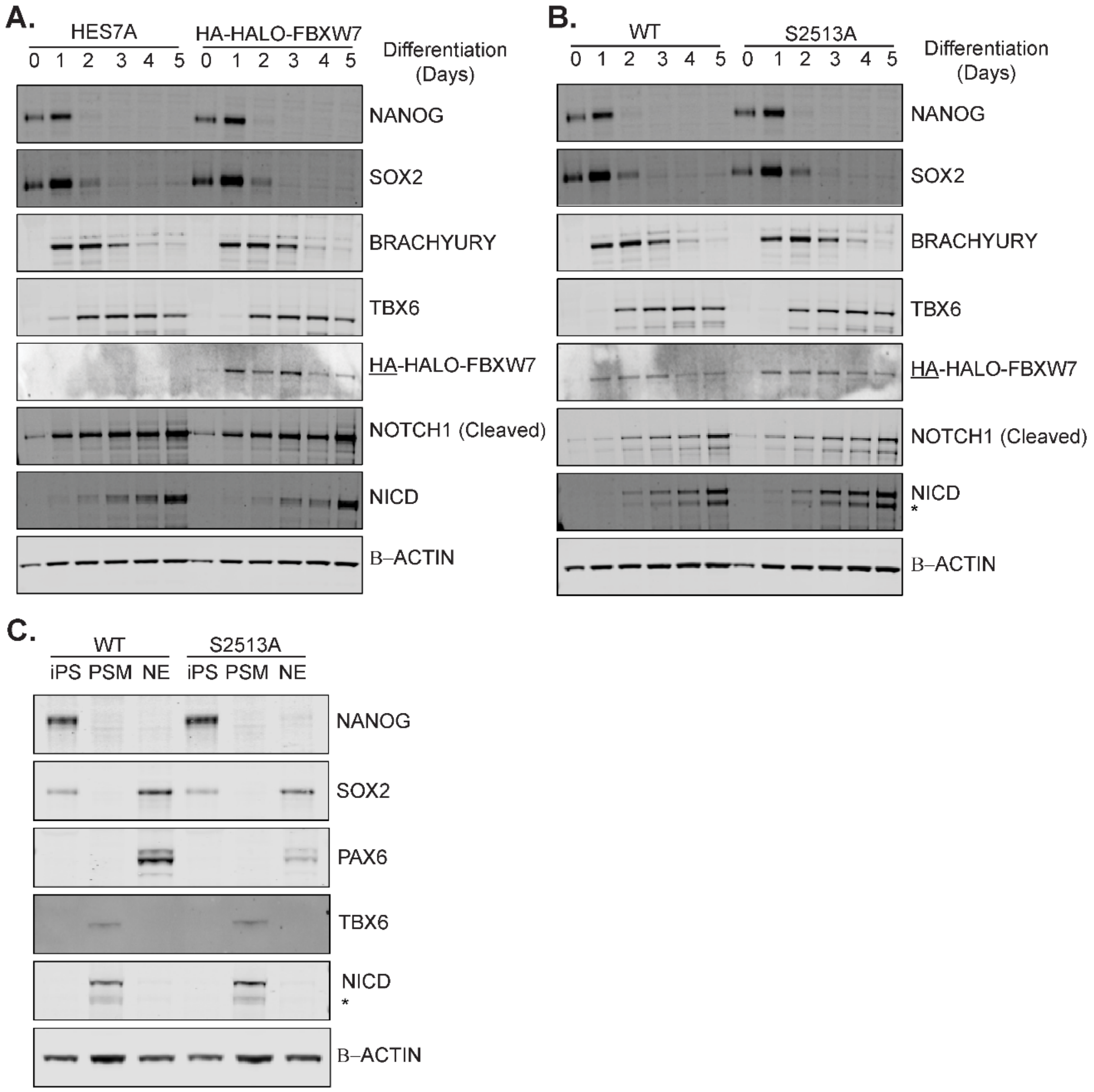

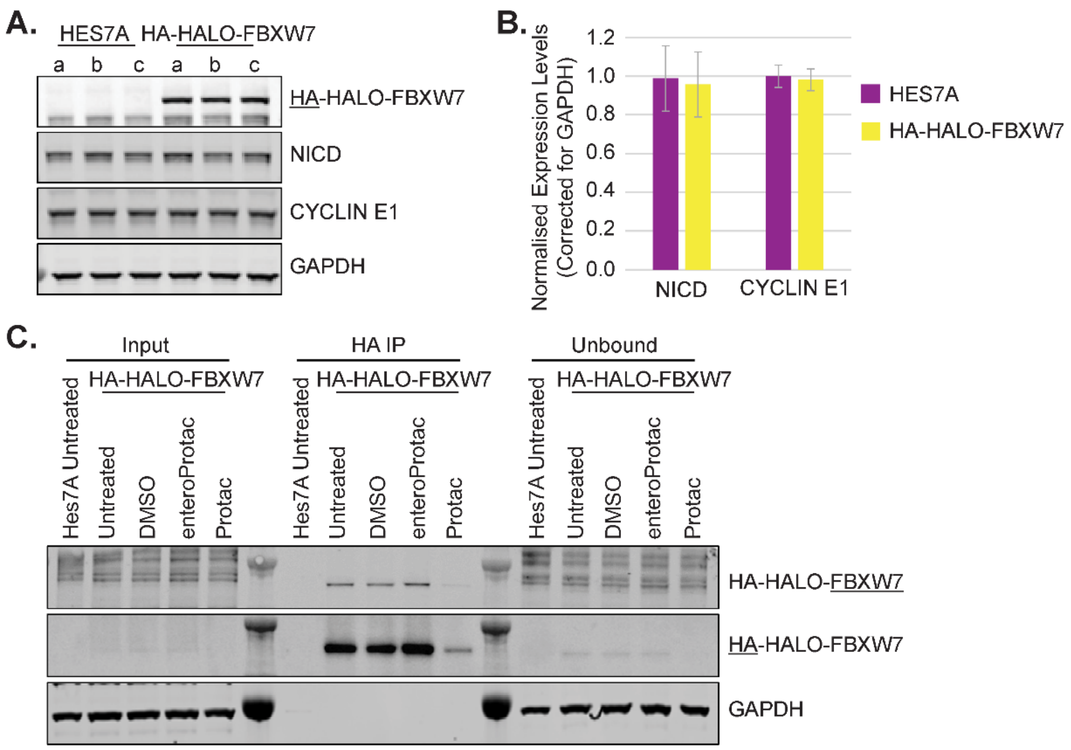

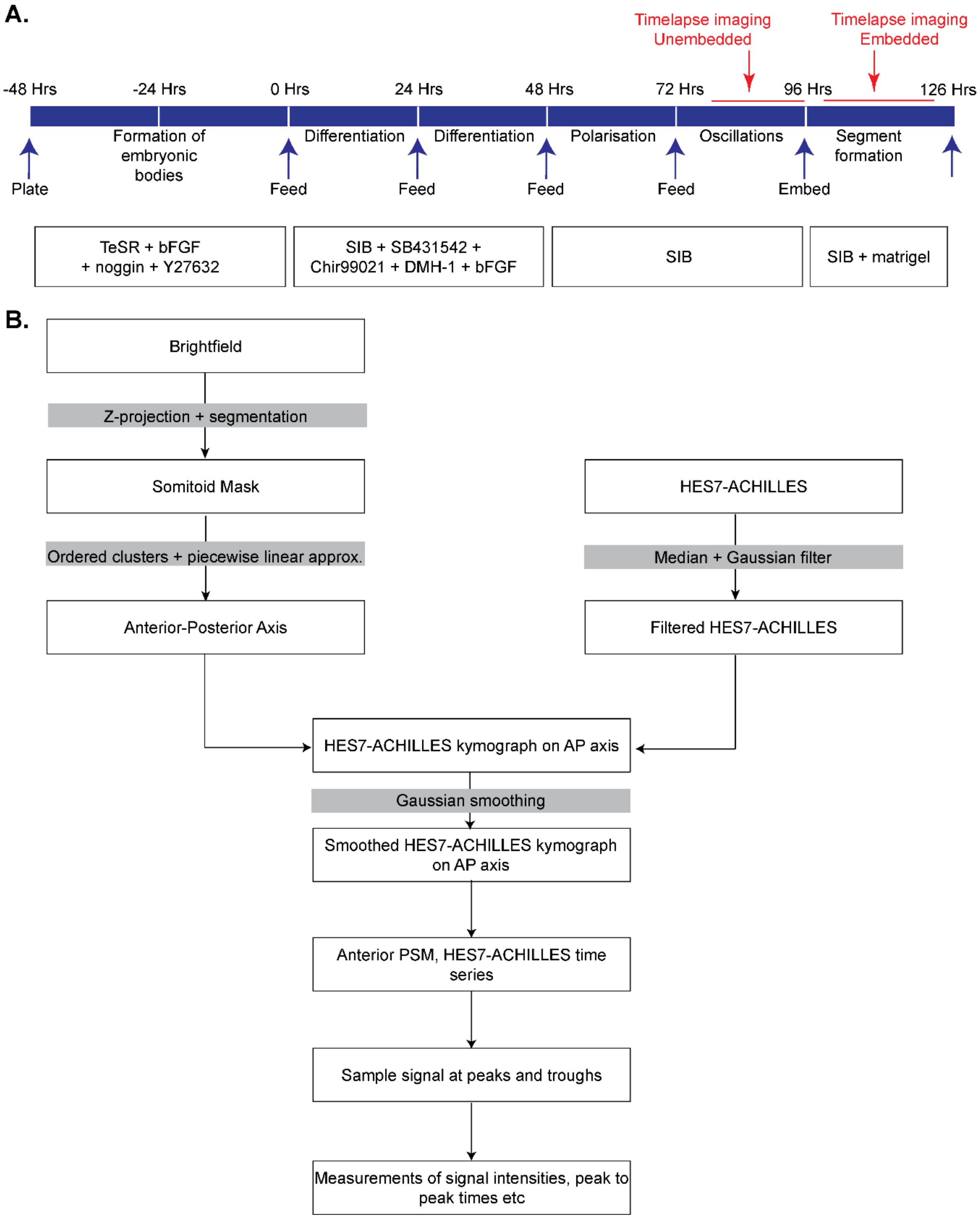

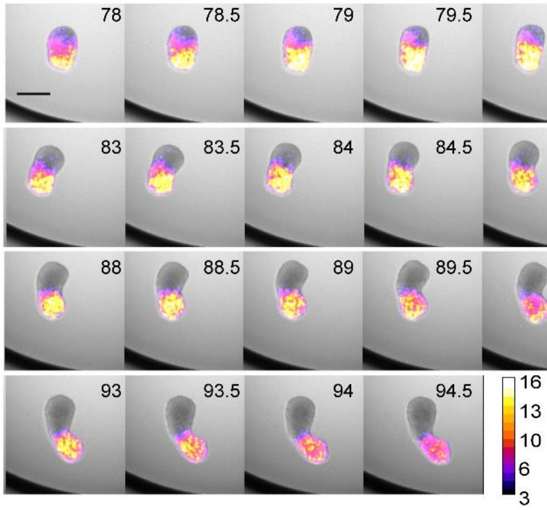

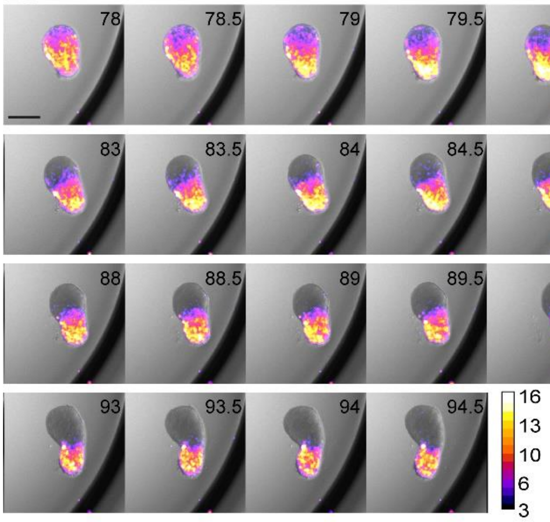

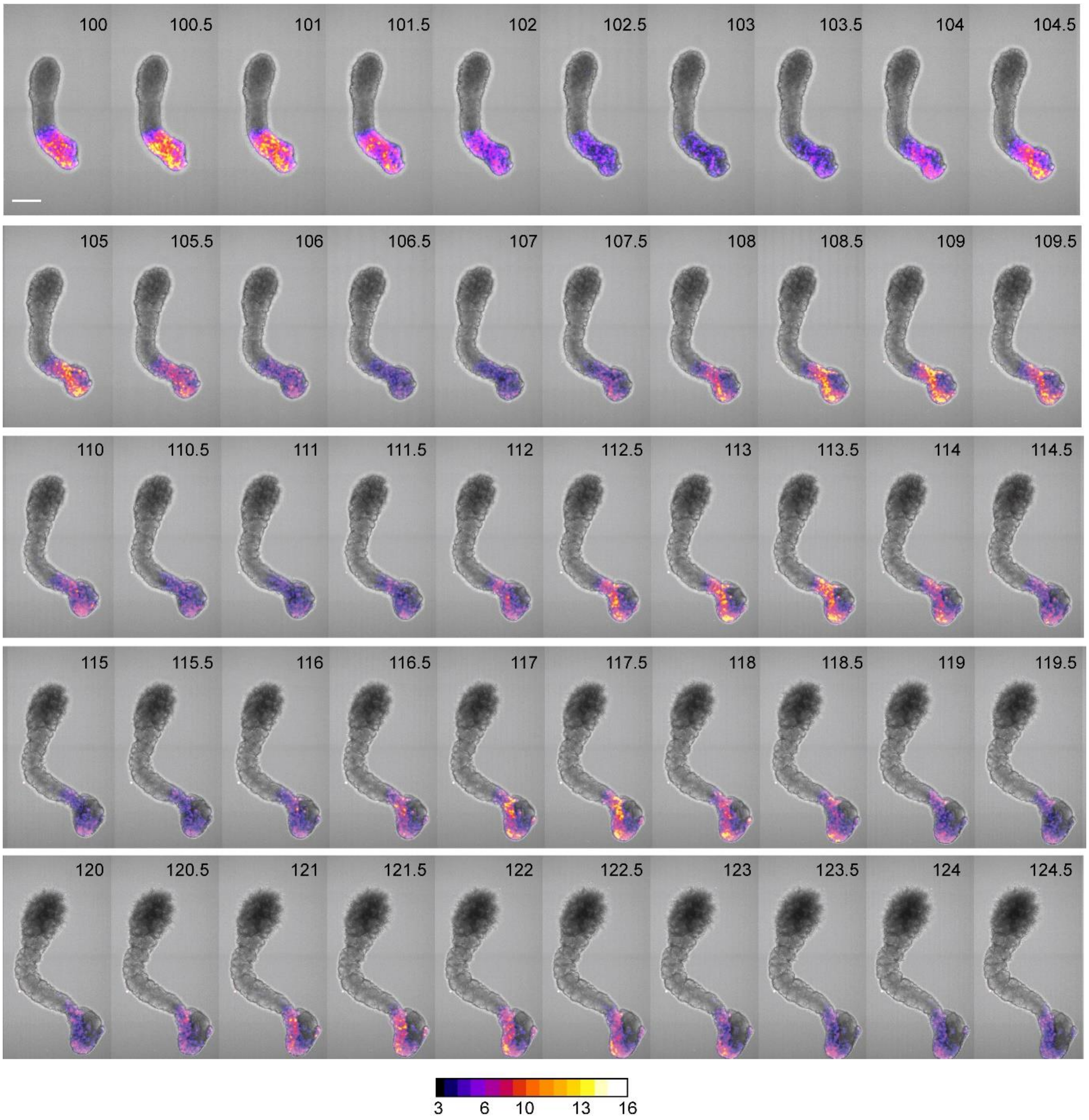

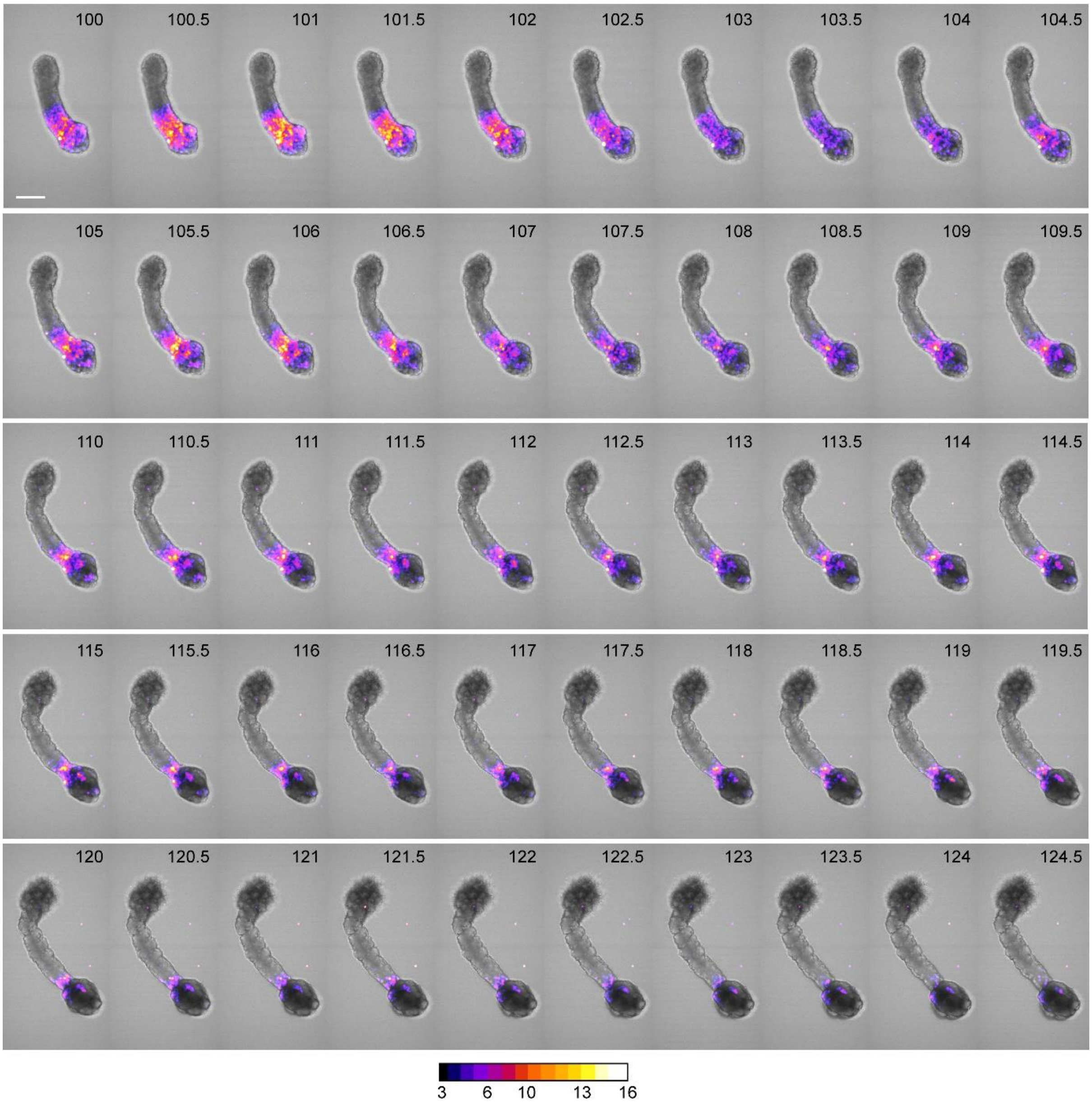

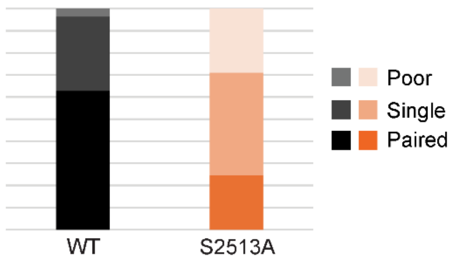
